# Thermoceptive predictions and prediction errors in the anterior insula

**DOI:** 10.1101/2024.10.11.617819

**Authors:** Birte Toussaint, Jakob Heinzle, Nicole Friedli, Nicole Jessica Zahnd, Elena Federici, Laura Köchli, Olivia Kate Harrison, Sandra Iglesias, Klaas Enno Stephan

## Abstract

Contemporary theories of interoception propose that the brain constructs a model of the body for predicting the states and allostatic needs of all organs, including the skin, and updates this model using prediction error signals. However, empirical tests of this proposal are scarce in humans. This computational neuroimaging study investigated the presence and location of thermoceptive predictions and prediction errors in the brain using probabilistic manipulations of skin temperature in a novel interoceptive learning paradigm. Using functional MRI in healthy volunteers, we found that a Bayesian model provided a better account of participants’ skin temperature predictions than a non-Bayesian model. Further, activity in a network including the anterior insula was associated with trial-wise predictions and precision-weighted prediction errors. Our findings provide further evidence that the anterior insula plays a key role in implementing the brain’s model of the body, and raise important questions about the structure of this model.

## Introduction

Perception is increasingly understood as a process of inference (1): confronted with noisy and ambiguous sensory signals, the brain is required to infer the latent (hidden) states of the world that give rise to its sensory inputs (2). This is frequently conceptualised as Bayesian inference, which rests on a generative model that combines a likelihood function (a probabilistic mapping from unobserved states to sensory data) with a prior belief about states of the world. Under this view, perception corresponds to the inversion of this model and is represented by a posterior (updated) belief (3–5).

Notably, in many statistical contexts, this process of perceptual inference or belief updating can be summarised as the interaction of two computational quantities: a prediction (based on the prior) and the prediction error (PE; how much the prediction deviates from the actual observation), weighted by precision (the relative uncertainty of predictions and sensory inputs). For this reason, computational theories of perception, such as predictive coding (2, 6), focus on how predictions and precision-weighted PE (pwPE) signals interact within neural circuits. In accordance with anatomical studies of sensory systems in cortex (7, 8), predictive coding (and related hierarchical Bayesian models of perception (9)) assume a processing hierarchy in which predictions and pwPEs are signalled across levels, leading to belief updates (inference) about hierarchically structured states of the world (e.g., spatial, temporal, or semantic hierarchies).

A large body of empirical evidence suggests that Bayesian inference constitutes a fundamental principle of exteroception, the perception of the external environment (for reviews, see (5, 10, 11)). In particular, many electrophysiological and neuroimaging studies have demonstrated temporal trajectories of prediction and pwPE signals that are consistent with Bayesian models, for example during visual (12–14), auditory (14, 15) and somatosensory (14, 16) paradigms.

Several prominent proposals (17–22) have extended this Bayesian framework to interoception, the perception of the physiological condition of the body (23). While these theories are influential and enjoy much popularity, there is as yet limited direct evidence that interoception indeed corresponds to a process of Bayesian inference. Rare exceptions include recent studies which applied a Bayesian learning model to behavioural responses to cardioceptive (heartbeat tapping (24)) and thermoceptive-nociceptive (thermal grill illusion (25)) tasks, respectively, providing results consistent with the notion that interoceptive processing follows Bayesian principles. However, beyond evidence from behavioural studies, an important step in testing Bayesian theories of interoception is the neurophysiological demonstration of trial-wise prediction and pwPE signals that conform to the proposed computational dynamics. To our knowledge, so far only one study has come close to providing empirical evidence of this sort. In this respiratory fMRI study, unexpected changes in breathing resistance during an associative learning paradigm evoked activations in the anterior insula that reflected trial-by-trial prediction certainty and PEs, providing initial evidence that the insula may contain a predictive model of bodily states (26). However, an important limitation of this study was that a Bayesian model was only marginally superior to a simpler associative learning (Rescorla-Wagner, RW) model in explaining behaviourally expressed interoceptive predictions; this result may have been due to the relatively small number of trials (a constraint of the breathing manipulation). As mandated by the pre-registered analysis plan, the simpler RW model was used for fMRI data analysis, demonstrating trial-wise activations of anterior insula by prediction certainty and (unweighted) PEs, but not allowing the authors to test directly for pwPEs.

Given this state of the literature on interoception, two major questions remain open: Is interoception of a Bayesian nature? And if so, what form does the corresponding generative model take, e.g. does it have a hierarchical structure, as proposed by predictive coding and related theories of hierarchical Bayesian inference?

In the present neuroimaging study, we tested for the presence of interoceptive predictions and pwPE signals in a task that involved probabilistic cues of imminent skin temperature changes. (It is worth highlighting that, while skin temperature was historically regarded as an aspect of exteroception (27), it is now widely accepted as an interoceptive function, as reflected in numerous reviews and theoretical analyses of interoception, e.g. (23, 20, 28–34)). Specifically, we developed a novel experimental paradigm that required active predictions of skin temperature changes (i.e., warming or cooling from a thermoneutral baseline) in a volatile learning environment (compare (12)). Following a pre-specified analysis plan (see below) and using formal model comparison, we contrasted a Bayesian learning model with a classical associative learning model (Rescorla-Wagner) to identify the most plausible mechanism at play, based on participants’ trial-wise predictions. The winning model was used to generate regressors of predictions and pwPEs for model-based fMRI analyses. We took two approaches to test different hypotheses: (i) analyses in pre-defined regions of interest (ROIs); and (ii) a more exploratory whole-brain analysis. Our primary ROI analysis focused on testing for the presence of thermoceptive predictions and pwPEs in the insula, a key node in contemporary theories of interoception. In short, our analyses showed that (i) behaviourally expressed predictions during our task were better explained by a Bayesian model than a non-Bayesian model, and that (ii) activity in the anterior insula reflected trial-wise predictions and pwPEs, as postulated by Bayesian inference.

## Results

Forty-four participants performed a skin temperature learning (STL) task while functional MR images and behavioural (button press) responses were acquired. In brief, the task required participants to learn the (changing) probabilistic associations between visual cues and skin temperature outcomes over 150 trials (see Figure 1). On each trial, one of two visual cues was presented, and participants had to predict (by button press) whether their skin would subsequently feel warm or cool.

**Figure 1.**
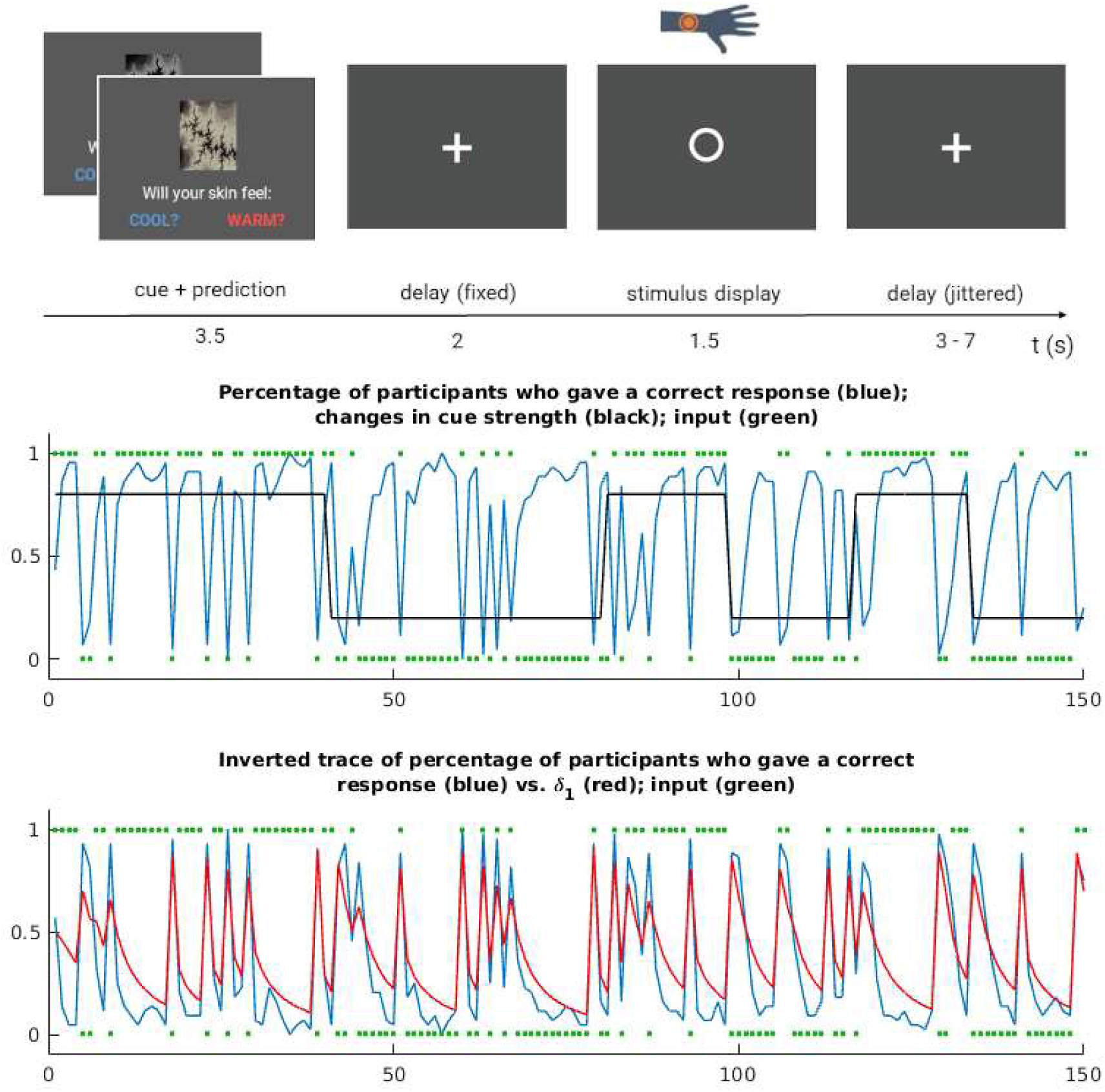
The skin temperature learning (STL) task for measuring dynamic learning of skin warming/cooling. *Top:* The structure of one trial. One of two cues was presented, based on which participants were asked to predict skin warming/cooling. After a fixed delay, the thermal stimulus was delivered while a circle appeared on the screen. The trial ended with a short delay of varying length. The task design had been optimised for statistical efficiency of the fMRI data analysis. *Middle:* The blue trace represents trial-wise correct responses as a percentage across the group of participants. The black trace represents the probability that one of the cues predicted a warm stimulus. The probability that the alternative cue predicted a warm stimulus is given by mirroring this trace (not shown). *Bottom:* Comparison between the inverted percentage trace from the panel above (i.e. trial-wise correct responses across participants; blue trace) and the absolute PE, *δ*1, averaged across participants (red trace). This qualitative comparison shows how the PE computed using the HGF resembles participants’ performance during the STL task.

The data collected during the STL task were used to address four research questions that were specified in a pre-registered analysis plan (which was electronically time stamped before any analyses were performed):

1. *Does a Bayesian model provide a better account of trial-wise predictions of skin temperature changes than a classical associative (non-Bayesian) learning model?*
2. *Does BOLD activity in the insula reflect interoceptive predictions and (precision-weighted) PEs regarding skin temperature?*
3. *Does BOLD activity in the dopaminergic midbrain reflect interoceptive predictions and (precision-weighted) PEs regarding skin temperature?*
4. *Are interoceptive predictions and (precision-weighted) PEs regarding skin temperature signalled within large-scale brain networks?*

(The inclusion of the precision weight in research questions 2-4 was contingent on the selection of the Bayesian learning model in research question 1.)

The results of the corresponding analyses are described in the following four subsections. While we show all results in the figures, our presentation in the main text focuses on activations (i.e. positive regression coefficients) by pwPEs and deactivations (i.e. negative regression coefficients) by predictions, in line with previous reports (26) and the role of prediction and pwPE terms in canonical Bayesian update equations (compare (9)).

### Behavioural data modelling

Using our pre-registered model space and random-effects Bayesian model selection (35, 36), we found that a Bayesian learning model (a hierarchical Gaussian filter (9, 37) (HGF) with two levels) provided a better account of participants’ behavioural data (i.e., button presses indicating the anticipation of skin warming/cooling) in the STL task than a simpler, non-Bayesian (Rescorla-Wagner) learning model (HGF: protected exceedance probability = 0.77; exceedance probability = 1.00; posterior probability = 0.75). In an additional validation step (not pre-specified in our analysis plan), we adopted the approach by Iglesias et al. (13), demonstrating that participants learned the probability structure of our paradigm in a way that closely mimicked the trajectory of pwPEs from our Bayesian model (see Figure 1).

The subsequent analyses of fMRI data addressed research questions 2-4 described above and were performed using model-based regressors of predictions (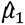) and pwPEs (*ɛ*_2_) based on the HGF. As participants’ predictions were categorical (i.e., warm or cool) and probabilities were coupled in the two learning trajectories (i.e., the probability of warming/cooling given one cue was 1 minus the probability of warming/cooling given the other cue), we considered the absolute value of fitted pwPE trajectories (this is equivalent to computing a separate trajectory for each categorical outcome; for details see the supplementary material in (12)). For notational simplicity, we refer to these pwPEs as *ɛ*_2_, rather than |ε_2_|, everywhere in this section.

### Prediction and pwPE activity in the insula

To test the hypothesis that the insula, a key node in Bayesian theories of interoception, signals predictions and pwPEs regarding skin temperature, we performed an ROI analysis based on a pre-specified mask of the bilateral insula. Peak-level, family-wise error (FWE)-corrected results (at a predefined significance threshold of *p* < 0.01; see *Methods*) are displayed in Table 1 and Figure 2. Overall predictions related to skin temperature outcome (i.e., averaged over the warming and cooling conditions) were associated with a significant deactivation in the bilateral dorsal anterior insula. Inspection of the separate contributions of the two underlying regressors to this effect revealed that deactivations associated with predictions about skin warming and cooling were located in overlapping clusters (see Figure S2 and Table S2). A smaller, but significant, positive correlation with overall predictions was observed in the right dorsal anterior-mid insula.

**Figure 2.**
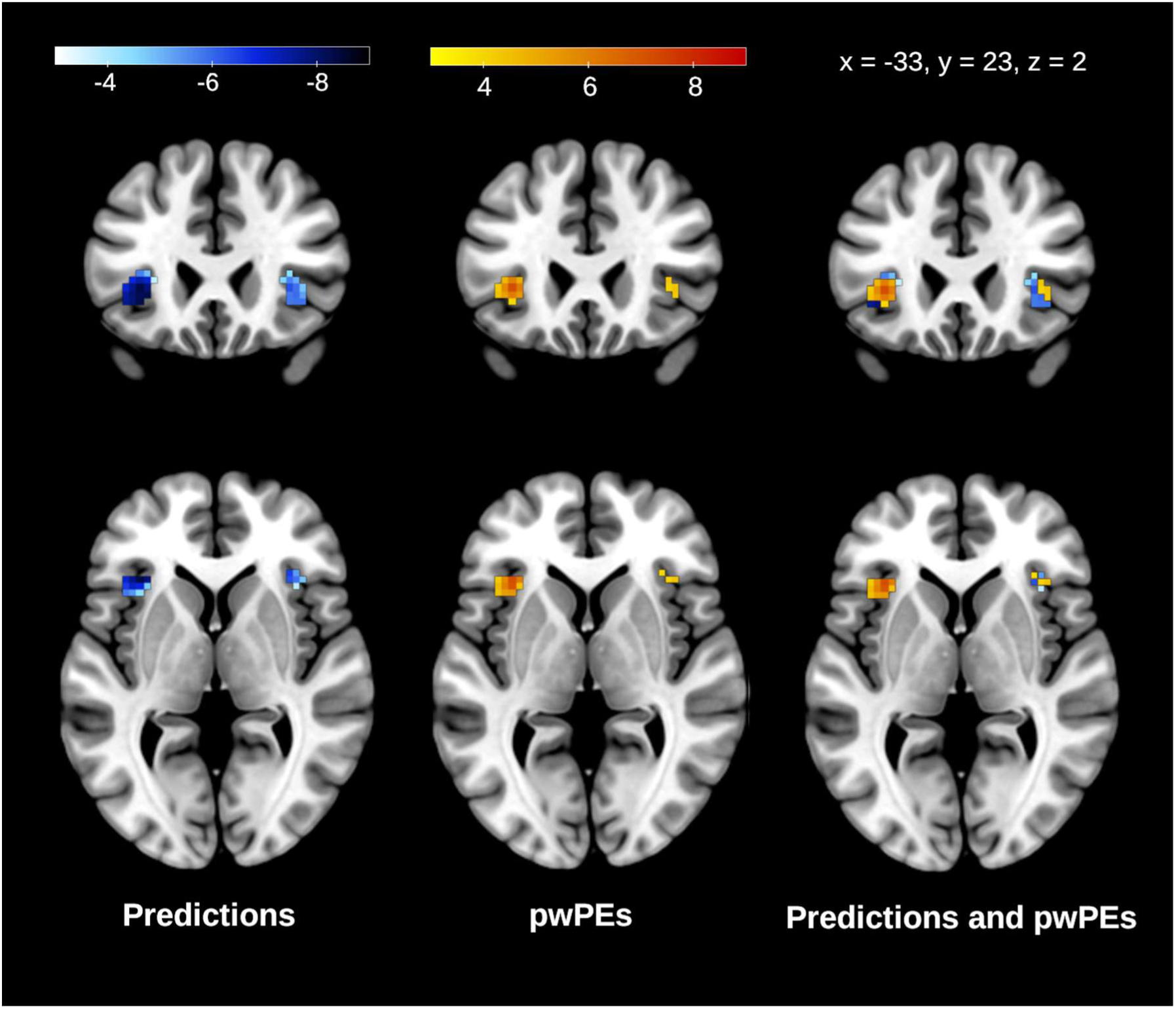
Subset of insular responses to predictions and precision-weighted prediction errors. *Left:* Deactivations associated with predictions about skin temperature outcome, 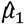, in the insular ROI. *Middle:* Activations associated with precision-weighted prediction errors about skin temperature outcome, *ɛ*_2_, in the insular ROI. *Right:* Activations with *ɛ*_2_ (warm colours) overlaid on deactivations with 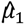 (cool colours). Activation maps are shown at a threshold of *p* < 0.01, peak-level FWE-corrected for multiple comparisons in the bilateral insula. (Please note that this figure only shows activations in our insular ROI. For further (de)activations, please see Figure 3).

**Table 1.** Insular responses to predictions and precision-weighted prediction errors. MNI coordinates and t-values for (de)activations associated with predictions, 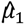, and activations associated with precision-weighted prediction errors, *ɛ*_2_, about skin temperature outcome. Note that activations have positive, deactivations negative t-values.

| **Predictions** |  |  |  |  |  |
| --- | --- | --- | --- | --- | --- |
|  | Hemisphere | x | y | z | t-score |
| Dorsal anterior-mid insula | R | 36 | 8 | 14 | 4.72 |
| Dorsal anterior insula | L | -33 | 23 | 2 | -8.44 |
| Dorsal anterior insula | L | -42 | 20 | -7 | -7.85 |
| Dorsal anterior insula | R | 33 | 26 | -1 | -7.29 |
| Ventral anterior insula | R | 42 | 20 | -10 | -5.25 |
| **Precision-weighted prediction errors** |  |  |  |  |  |
| Dorsal anterior insula | L | -33 | 23 | 2 | 7.49 |
| Dorsal anterior insula | R | 39 | 23 | 2 | 5.20 |
| All results: $p < 0.01$ peak-level FWE-corrected for multiple comparisons in the bilateral insula. | | | | | |

Overall pwPEs related to skin temperature outcome (i.e., averaged over the warming and cooling conditions) were associated with an activation of the bilateral dorsal anterior insula, with pwPEs related to skin warming and cooling located in neighbouring clusters (see Figure S3 and Table S3). These activations by pwPEs were nearly identical in location and extent to the deactivations by predictions (see right column of Figure 2).

### Prediction activity in the midbrain

In a next step, we tested the hypothesis that the dopamine system is recruited in interoceptive signalling, using an ROI analysis of the dopaminergic midbrain (i.e., the substantia nigra, SN, and ventral tegmental area, VTA). This analysis was motivated by findings from an earlier study that employed a similar associative learning task in the exteroceptive domain and showed that dopaminergic midbrain activity reflects pwPEs regarding purely sensory inputs (12). The role of dopamine in signalling sensory PEs unrelated to reward has since been corroborated in other work (e.g., (38–42)). Here, we asked whether midbrain responses might also reflect interoceptive prediction and pwPE signals. Peak-level FWE-corrected results (at a predefined significance threshold of *p* < 0.01; see *Methods*) are presented in Table 2 and Figure S1. The dopaminergic midbrain demonstrated a significant deactivation with overall predictions of skin temperature outcome. We also observed an activation with pwPEs (*p* = 0.029, FWE-corrected) but this effect did not survive correction across the number of statistical tests (which required *p* < 0.01 FWE-corrected for significance; see *Methods*).

**Table 2.** Midbrain responses to predictions. MNI coordinates and t-values for deactivations associated with predictions about skin temperature outcome, 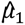.

| x | y | z | t-score |
| --- | --- | --- | --- |
| 3 | -22 | -22 | -4.78 |
| -3 | -19 | -22 | -4.09 |
| All results: $p < 0.01$ peak-level FWE-corrected for multiple comparisons in the SN/VTA. | | | |

### Whole-brain analysis of predictions and pwPEs

We complemented our focused ROI analyses with a whole-brain analysis to identify potentially distributed brain networks involved in processing predictions and pwPEs related to skin temperature. Key results are displayed in Table 3 and Figure 3, with additional figures and tables provided in the supplementary material. As pre-specified in our analysis plan, in this whole-brain analysis we tested for significance at the cluster level; therefore, the locations of peak activations within clusters are indicated here and in the results tables for descriptive purposes only. Additionally, for completeness, we report cluster peaks that survived correction at the peak level (*p* < 0.025, FWE-corrected), but emphasise that peak-level statistics were not used to determine significance. Overall (i.e., averaged over the warming and cooling conditions) predictions of skin temperature outcome were associated with widespread deactivations in 7 significant clusters (see Figure 3). The peaks of these clusters were located in the left dorsolateral prefrontal cortex, left and right inferior parietal cortex, left ventral caudate, left and right cerebellum, and right precuneus. A qualitative comparison (see Figure S4 and Table S4) shows a similar pattern of deactivations with predictions of skin warming and skin cooling, respectively. Significant activations with predictions are also displayed in Table 3 and Figure 3.

**Figure 3.**
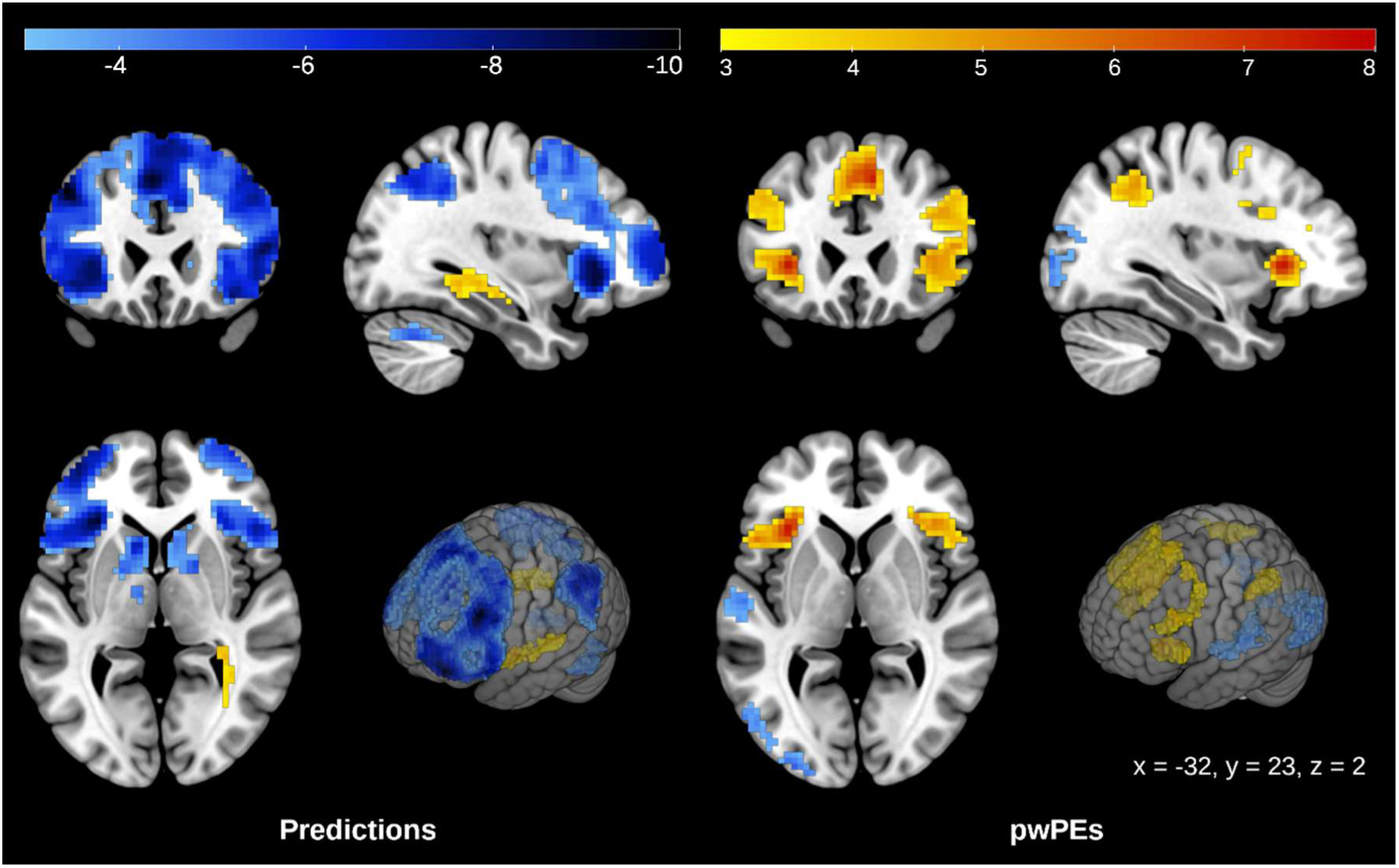
Whole-brain results: (De)activation with predictions and precision-weighted prediction errors. *Left:* (De)activations with predictions about skin temperature outcome, 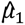. *Right:* (De)activations with precision-weighted prediction errors about skin temperature outcome, *ɛ*_2_. Activations are shown at *p* < 0.025 cluster-level FWE-corrected across the whole brain (with a cluster-defining threshold of *p* < 0.001).

**Table 3.** Whole-brain results: Modulation of neural activity by predictions and precision-weighted prediction errors. Extent of clusters (number of significant voxels) for (de)activations associated with predictions, 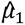, and precision-weighted prediction errors, *ɛ*_2_, about skin temperature outcome. MNI coordinates and t-values are listed for peaks within significant clusters. Note that activations have positive, deactivations negative t-values. (De)activations for which *p* < 0.025 at the peak level are indicated with an asterisk.

| Predictions |  |  |  |  |  |  |
| --- | --- | --- | --- | --- | --- | --- |
| Extent | Anatomical label (Brainnetome atlas) | Hemisphere | x | y | z | t-score |
| 123 | Rostral hippocampus | L | -27 | -13 | -16 | 5.54* |
| 196 | Precentral gyrus | R | 39 | 8 | 14 | 5.22 |
| 163 | White matter/cerebrospinal fluid | R | 30 | -40 | 11 | 4.97 |
| 5922 | Middle frontal gyrus | L | -45 | 23 | 38 | -9.95* |
|  | Orbital gyrus | L | -30 | 29 | -4 | -9.94* |
|  | Superior frontal gyrus | L | -3 | 23 | 44 | -9.30* |
| 723 | Intraparietal sulcus (hIP3) | L | -42 | -52 | 47 | -8.27* |
|  | Supramarginal gyrus (PFm) | L | -54 | -46 | 47 | -7.63* |
|  | Intraparietal sulcus (hIP3) | L | -39 | -58 | 41 | -7.47* |
| 857 | Intraparietal sulcus (hIP3) | R | 48 | -61 | 38 | -7.54* |
|  | Supramarginal gyrus (hIP3) | R | 51 | -49 | 38 | -7.34* |
|  | Intraparietal sulcus (hIP3) | R | 39 | -61 | 47 | -7.10* |
| 320 | Ventral caudate | L | -9 | 14 | 2 | -7.06* |
|  | Ventral caudate | L | -12 | 2 | -4 | -6.52* |
|  | Globus pallidus | R | 15 | 2 | 5 | -5.90* |
| 85 | Cerebellum | R | 36 | -58 | -34 | -6.52* |
| 222 | Precuneus | R | 6 | -64 | 47 | -6.26* |
| 110 | Cerebellum | L | -30 | -61 | -34 | -5.56* |
| Precision-weighted prediction errors |  |  |  |  |  |  |
| 168 | Dorsal anterior insula | L | -33 | 23 | 2 | 7.50* |
|  | Inferior frontal gyrus | L | -48 | 17 | 2 | 5.67* |
| 581 | Superior frontal gyrus | R | 9 | 23 | 44 | 7.40* |
|  | Superior frontal gyrus | R | 3 | 32 | 47 | 6.93* |
|  | Superior frontal gyrus | L | -6 | 14 | 53 | 6.84* |
| 712 | Inferior frontal gyrus | R | 48 | 20 | 8 | 5.55* |
| 228 | Inferior frontal junction | L | -42 | 17 | 29 | 5.33 |
| 144 | Intraparietal sulcus (hIP3) | L | -36 | -49 | 41 | 5.13 |
| 112 | Intraparietal sulcus (hIP3) | R | 36 | -55 | 41 | 4.98 |
| 355 | Occipital pole | L | -30 | -94 | 5 | -5.71* |
| 230 | Ventromedial parietooccipital sulcus | L | -6 | -64 | 17 | -5.01 |
| 205 | Superior temporal gyrus (Te1.0 & Te1.2) | L | -54 | -13 | 2 | -4.67 |

All clusters: $p < 0.025$ , cluster-level FWE-corrected for multiple comparisons across the whole brain (cluster-defining threshold: $p < 0.001$ ).
\* $p < 0.025$ , peak-level FWE-corrected for multiple comparisons across the whole brain.

The significant activation map for overall pwPEs related to skin temperature outcome (i.e., averaged over the warming and cooling conditions) demonstrated a similar pattern as that observed for deactivations associated with predictions (see Figure 3). The peaks of the 6 significant clusters were located in the left anterior insula, right dorsolateral prefrontal cortex, right ventrolateral prefrontal cortex, left inferior frontal junction, and left and right inferior parietal cortex. As with predictions, pwPEs related to warming and cooling displayed similar activation patterns (see Figure S5 and Table S5). All significant (de)activations with pwPEs are reported in Table 3 and displayed in Figure 3.

## Discussion

The concept of perception as Bayesian inference has become highly popular. For exteroceptive modalities, such as vision, this notion is backed by extensive empirical evidence (5, 10). More recently, Bayesian explanations have also been applied to interoception (17–22). So far, however, there is only limited direct evidence from empirical studies that interoception is grounded in Bayesian inference (24–26) (see *Introduction* for more details).

In contrast to the scarcity of empirical evidence, theoretical proposals detail how interoceptive Bayesian inference processes may be implemented in the brain (20, 22, 43). These proposals typically assume hierarchically structured circuits, spanning multiple areas of the insula in particular, which implement a form of hierarchical Bayesian inference, such as predictive coding (2). In analogy to previously established concepts for exteroception (e.g. vision), hierarchical Bayesian concepts of interoception build on anatomically defined hierarchical relationships between cortical areas (20, 22, 43). However, unlike the anatomical hierarchies underlying exteroception (which derive from laminar patterns of cortical connections), the structural hierarchies proposed for interoception rest on indirect anatomical evidence (i.e. regional differences in cytoarchitecture (44)) due to a lack of lamina-level tract tracing data in primates for key interoceptive regions like the insula. Furthermore, direct functional evidence that a hierarchical form of Bayesian inference underlies interoception is lacking so far.

Despite the theoretical plausibility and general appeal of (hierarchical) Bayesian theories of interoception, the relative lack of anatomical and functional evidence means that two major questions remain open: (i) whether interoception corresponds to Bayesian inference at all, and if so, (ii) whether the corresponding generative model has a hierarchical structure, as assumed by predictive coding and related theories of hierarchical Bayesian inference.

In this paper, we presented results from a computational neuroimaging study that contribute towards clarifying these questions. Our results support the general notion that interoception rests on Bayesian inference: random-effects Bayesian model selection showed that a Bayesian model (a two-level HGF) provided a better explanation of participants’ behaviour than a non-Bayesian associative learning model (Rescorla-Wagner). Further, our fMRI analyses demonstrated that activity in the anterior insula showed a hallmark of Bayesian inference, i.e. trial-by-trial pwPEs and predictions.

By contrast, concerning the question of how Bayesian inference may be implemented in the brain, it is not straightforward to reconcile our results with the notion that interoception rests on a Bayesian inference hierarchy in the insula. This is because we observed pwPE responses only within the anterior insula, which constitutes a "high-level” area within proposed interoceptive hierarchies (20, 22), but not in putatively "low-level” areas such as the posterior insula. This finding is not consistent with hierarchical Bayesian schemes like predictive coding or hierarchical Gaussian filtering where, upon the arrival of sensory input at the bottom of the hierarchy, pwPEs cascade up the hierarchy until they reach a level where beliefs no longer require updating. In other words, in these hierarchical schemes it is possible to observe that sensory inputs evoke pwPEs in lower but not higher areas, while the opposite response pattern does not occur. It is noteworthy that a previous study reporting trial-wise prediction error activations during interoception (using breathing resistance stimuli (26)) found activations by respiratory PEs and predictions in the bilateral anterior insula that correspond very closely to activations by thermoceptive pwPEs and predictions in our study – although it should be kept in mind that the computational models differed between the two studies.

At this point one may ask why the computational model we applied to fMRI data only contains a single type of pwPE, not multiple hierarchically related ones. It is worth highlighting that a central challenge for hierarchical Bayesian concepts of perception is to define the exact nature of the hierarchy, i.e. to specify the quantities that are represented at each hierarchical level. For example, for visual and auditory hierarchies, concrete suggestions exist (e.g. spatial, temporal, and/or semantic hierarchies), some of which have also been implemented in hierarchical models applied to empirical data (e.g. in the visual (6, 45, 46) and auditory (47–51) domains). By contrast, we are not aware of any concrete proposals concerning the representational nature of a computational hierarchy for interoception. In the absence of an exact concept of the hierarchy, one strategy for delineating the approximate extent of a perceptual inference hierarchy is to map the activations elicited by the occurrence of surprising sensory events (such as "oddballs" or "deviants"; e.g. (52–56)) or, more quantitatively, low-level PEs about sensory events (e.g. (15, 57)). (The latter is the strategy used in our fMRI analysis.) This approach is motivated by the process already described above, i.e. in standard implementations of predictive coding, pwPEs propagate across levels and are correlated (since a PE at a higher level can only be triggered by the arrival of a PE from a lower level). Put differently, when a surprising sensory event occurs, all levels of the inference hierarchy that require belief updating should show activation.

Overall, our findings thus support the notion that interoception corresponds to a process of Bayesian inference but cannot be easily reconciled with contemporary interoceptive predictive coding proposals. This raises the question how this constellation of results might be explained. In the following we consider two possibilities:

First, our failure to find pwPEs in posterior regions of the insula – which are thought to represent low levels of the interoceptive inference hierarchy (20, 22) – questions whether these areas are part of the inference hierarchy for interoception, or whether they might rather serve to represent sensory inputs from the body, including temperature information, and relay this information to anterior areas in which the inference process takes place. This scenario is compatible with concepts that assign representation of objective temperature to posterior insula and of subjective (i.e. perceived) temperature to anterior insula (58, 59). This possibility – i.e. posterior insular areas serving to represent and relay sensory inputs from the body, and anterior insular areas acting as a “comparator” of interoceptive information – has also featured in previous proposals (17) (see also (60)). While our results are not easy to reconcile with an *insular* Bayesian inference hierarchy, the overlap of prediction and pwPE activity in several cortical areas (see Figure 3) is compatible with the possibility that the anterior insula lies at the lowest level of a cortical hierarchy implementing interoceptive Bayesian inference. Our post-hoc finding that temperature changes activate different regions of the insula (see Figure S6 and Table S6) than temperature-related prediction and pwPE responses is consistent with this interpretation. Moreover, a similar functional dissociation was observed in a neuroimaging study on pain perception, which found that stimulus intensity was encoded in the posterior insula, while pain expectation and error signals resulting from unexpected pain were encoded in the anterior insula (61).

A second possibility is that the brain might switch between different implementations of Bayesian inference, depending on task demands (62) (e.g. whether adequate task performance can only be achieved with a complex and spatially distributed hierarchical model or whether a more local model might suffice). This possibility is supported by results from a parallel experiment (only described in a PhD thesis so far) (63) which used equivalent thermal stimuli to those in the current study, and also varied their predictability. However, this parallel experiment did not require any behavioural responses to the stimuli – nor did it include any cues or provide prior structural information about the paradigm, as in the current study, therefore posing a more difficult (less constrained) inference problem. In this implicit paradigm, prediction error responses were found throughout the entire insula, from posterior to anterior areas (63). This result may reflect the fact that the inference problem was more challenging than in the current study, creating greater model uncertainty, and potentially leading to the deployment of more complex and diverse implementations of Bayesian inference. Interestingly, dissociations between explicit and implicit forms of Bayesian inference have also been reported in behavioural studies (64–66).

These explanations are tentative but serve to outline how our current findings might be reconcilable with classical predictive coding accounts of interoception. We emphasise that these explanations are highly speculative at this point and would have to be tested in suitably designed future studies.

Our study has several strengths and limitations that should be kept in mind when interpreting its results. Beginning with strengths, it transferred a previously established learning paradigm from the exteroceptive domain (12) to interoception, using thermostimulation of the skin. This stimulation method allows for more trials per experimental session than has been possible with structurally comparable respiratory investigations of interoception (26). Furthermore, several measures were taken to ensure that the analyses were robust and protected against (cognitive) biases. For example, all analyses were based on a preregistered analysis plan, including the prior specification of models to be tested. Furthermore, priors for the parameters of all models were derived from an independent data set, thus creating a level playing field for model comparison (which is sensitive to the choice of priors).

Concerning limitations, the results from our random-effects Bayesian model selection procedure were not very strong, implying a degree of heterogeneity in the sample we studied. Most importantly, the voxel-wise general linear model analysis is frequentist in nature and does not allow for accepting the null hypothesis. In other words, we cannot quantify our certainty about the absence of pwPE responses in posterior regions of the insula; we can only say that we failed to reject the null hypothesis (even when examining responses at a lenient threshold of *p* < 0.001 uncorrected, no significant pwPE signals were observed).

In summary, our findings support the general notion that interoception draws on principles of Bayesian inference. More specifically, they extend previous initial evidence that the anterior insula plays a key role in implementing a generative model of sensory inputs from the body (26). Our results raise questions about the (hierarchical or non-hierarchical) structure of this generative model – and whether it has a constant form across all contexts – that require further scrutiny in future studies. Given the important role of interoceptive processing in mental health conditions, this line of research has the potential to offer a mechanistic understanding of underlying pathologies (29, 67).

## Methods

### Participants

Overall, our study included 64 healthy, right-handed participants in two datasets. The MRI data set consisted of N = 44 individuals (30 females, aged (mean ± SD) 24.9 ± 4.2 years), while an independent behavioural data set (that was used to derive priors for the model-based fMRI analysis) included N = 20 participants (11 females, aged (mean ± SD) 25.9 ± 4.6 years). Participants were pre-screened online to ensure they did not display clinically significant levels of depression (indicated by a score above 16 in the Center for Epidemiologic Studies Depression Scale (68)) or central sensitisation syndrome (indicated by a score above 40 in section A of the Central Sensitization Inventory (69)). Further exclusion criteria are detailed in the supporting information. The research protocol was approved by the Cantonal Ethics Committee Zurich (ethics approval BASEC-No. 2018-02386). All participants provided their written informed consent and were compensated financially for their participation in the study.

### Sample size

In our analysis plan, we had specified a Sequential Bayes Factor (70) (SBF) design for our MRI data set. However, due to technical issues with the thermostimulation system, we were forced to abandon our predefined SBF design and instead collected a final sample of 44 data sets. Please refer to the supporting information for further details. For the estimation of priors in the computational models, we collected an independent behavioural data set, for which we had pre-specified a fixed sample size of N = 20 in our analysis plan.

### Thermal stimulation

The experimental stimuli consisted of warm (39 °C) and cool (27 °C) temperature pulses, applied with an MRI-compatible contact thermal stimulator (Pathway, Medoc Ltd., Ramat Yishai, Israel). The thermode (type CHEPS, 27 mm diameter) was secured approximately 10 cm from the wrist along the participant’s left dorsal forearm. A continuous baseline temperature of 32 °C was maintained between warm and cool temperature pulses. The rates of heating and cooling from baseline were set to 28 °C/s and 20 °C/s, respectively. After each warm or cool pulse, the temperature was actively returned to baseline at a rate of 40 °C/s. During each warm or cool temperature pulse the target temperature was applied for 0.7 s.

### Magnetic resonance imaging

The experiment was conducted on a 3 Tesla Philips Ingenia scanner (Philips Healthcare, Best, The Netherlands), using a 32-channel head coil. For functional imaging, 30 slices were acquired in ascending order using a fast (TR = 1 s) T_2_*-weighted echo-planar imaging sequence (voxel size = 3 x 3 x 3 mm^3^, inter-slice gap = 0.6 mm, field of view = 220 x 220 x 107.4 mm^3^, TE = 31 ms, flip angle = 70°, SENSE factor = 2.2, multi-band factor = 2). The run lasted approximately 30 minutes (1860 volumes) and began with the acquisition of ten dummy volumes to account for magnetisation saturation effects. A second-order shim (automatically adjusted by the scanner) was applied to reduce field inhomogeneities. A T_1_-weighted MPRAGE structural image was collected using a 3D T1-Inversion Recovery Turbo-Field-Echo sequence (voxel size = 1 x 1 x 1 mm^3^, field of view = 256 x 256 x 200 mm^3^, TR = 8.3 ms, TE = 3.85 ms, flip angle = 8°, shot TR = 2800 ms, inversion time = 982 ms, SENSE factor = 1.4). The structural scan lasted 5.3 minutes. Respiratory, cardiac and pulse activity was recorded simultaneously with EPI acquisition using a pneumatic breathing belt, a 4-electrode electrocardiogram, and a pulse oximeter (attached to the participant’s left index finger). Behavioural data was measured using a response box (Current Designs, with Psychophysics Toolbox Version 3 (PTB-3), version 3.0.15, http://psychtoolbox.org/) on which participants responded by pressing one of two buttons (right index and middle finger).

### Experimental protocol

After setting the participant up in the scanner (or in the behavioural laboratory), with the thermode attached to the left dorsal forearm, a continuous stimulation at the baseline temperature (32 °C) was applied for 5 minutes to ensure that the skin in contact with the thermode adapted completely to this temperature. Next, we presented the experimental stimuli and performed several control checks to ensure that the participant could clearly identify the stimuli and did not experience any confounding perceptual effects (such as pain). These control checks are described in the supporting information.

The participant then performed the skin temperature learning (STL) task (see Figure 1). In this paradigm, the participant was required to learn the probabilistic association of visual cues (presented using PTB-3) with skin temperature outcomes. On each trial, one of two greyscale images of fractals was presented for 3.5 seconds, during which time the participant had to predict (by button press) whether their skin would subsequently feel warm or cool. Participants were instructed to respond as accurately and quickly as possible, and were incentivised to do so with a performance bonus based on the reaction time for correctly predicted skin temperature in five randomly selected trials. The order of response options (warm and cool) and the cue-outcome pairs (i.e., which of the two fractals was more predictive of skin warming/cooling) were counterbalanced across participants. After the participant’s prediction (or after the display of a time-out screen, if no prediction was made within 3.5 seconds), there was a 2-second delay before the thermal stimulus was applied. Stimulus onset asynchrony (SOA) lengths were jittered between 10 and 14 seconds. (The task design was optimised for efficiency.) We instructed participants explicitly that the association strength for a particular cue-outcome pair could change over the course of the experiment, and that the probabilities of the cues were coupled (i.e., the probability of outcome A given one cue was 1 minus the probability of outcome A given the other cue). The task consisted of 150 trials divided into a stable phase (2 blocks of 40 trials) and a volatile phase (2 blocks of 18 trials and 2 blocks of 17 trials).

After completion of the STL task, the participant performed a further control task that allowed us to check for habituation effects (described in the supporting information). Finally, we performed a 5-minute structural scan. The participant completed a debriefing questionnaire outside the scanner.

### Exclusion and replacement of participants

Prior to data collection, we had defined criteria in our analysis plan for excluding and replacing participants due to data issues in the categories of thermosensation, behavioural responses, and MRI data quality. These criteria are detailed in the supporting information.

### Data analysis

The analysis steps described below were pre-specified in an analysis plan (https://gitlab.ethz.ch/tnu/analysis-plans/toussaintetal_compi_stl) that was finalised and time stamped before any statistical analyses (i.e. hypothesis testing or inference) were conducted. Preprocessing of MRI data was performed concurrently with data collection to determine whether data sets needed to be replaced. No other inspections of the data were performed before the analysis plan was uploaded.

### Preprocessing

MRI data were preprocessed using SPM12 (http://www.fil.ion.ucl.ac.uk/spm). Each participant’s functional images were corrected for slice timing and realigned to the mean functional image. The anatomical image was processed using the unified segmentation procedure (71). The mean functional image was coregistered to the resulting bias-corrected anatomical image, and the derived normalisation fields were applied to all functional images. The functional images were resampled to 3 x 3 x 3 mm^3^ voxels and smoothed using a 6 mm full width at half maximum Gaussian kernel.

In our statistical analyses, we accounted for head motion and physiological confounds by including them as nuisance regressors in the GLM. For details, please see the supporting information.

### Computational modelling of behaviour

An overview of the steps in our computational modelling pipeline is shown in Figure S7.

To test whether Bayesian inference provides a plausible mechanistic explanation of learning in the STL task, we formally compared the explanatory power of a Bayesian learning model with a simpler, non-Bayesian alternative. Below, we first present the learning models (including the specification of prior parameters), followed by a description of the model comparison procedure.

The models were implemented within the same computational framework using the HGF toolbox (9, 37) in the TAPAS software. This framework consists of two components: a perceptual model that maps environmental causes to sensory inputs, and a response model that maps the (inferred) environmental causes to a participant’s observed behavioural data (72).

### Response model

For the models considered in our analysis, we specified the same probabilistic response model, namely the unit-square sigmoid (37):

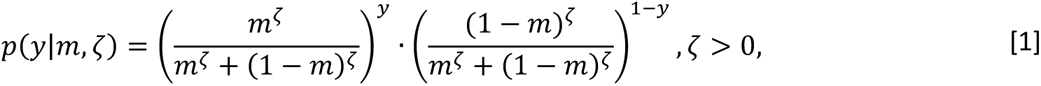

where *y* represents a participant’s trial-wise responses, *m* represents a participant’s beliefs about the association between visual cues and thermal stimuli, and *ζ* is a subject-specific free parameter encoding inverse response noise. (Note that *y* and *m* are vectors containing one element for each trial.) In our setting, *m*^(*k*)^ is the predictive probability that cue 1 is followed by a warm stimulus (and cue 2 by a cool stimulus) on trial (*k*). Thus, this model maps participants’ beliefs onto the probabilities *p*(*y*^(*k*)^ = 1) and *p*(*y*^(*k*)^ = 0) that the participant chose response 1 (i.e., predicted a warm stimulus when cue 1 was presented) or 0 (i.e., predicted a cool stimulus when cue 1 was presented), respectively, using a sigmoid function. Importantly, this (arbitrary) 1/0 coding of cue-outcome contingencies did not influence results in our study as we recoded the computational trajectories obtained in contingency space into temperature space (see *Model-based regressors* and the supplementary material in (12)).

### Perceptual model 1: Hierarchical Gaussian Filter

The Hierarchical Gaussian Filter (9) (HGF) is a generic Bayesian model that is frequently employed in studies of learning under uncertainty (for example, (12, 15, 26, 73–75)). The HGF maps an agent’s beliefs about the causes of sensory inputs onto his/her observed behavioural responses. In the supporting information, we provide a brief outline of the HGF implementation in our study; a detailed description of the inversion scheme and the specific update equations can be found in the papers that introduced the HGF (9, 37). We used the eHGF functions introduced in version 6.0 of the HGF toolbox with the built-in multi-start approach for model inversion.

The HGF can be used to model learning at multiple levels. As specified in our analysis plan, we did not make an *a priori* assumption about the hierarchical depth of our Bayesian model, but compared the suitability of HGF implementations with two and three levels by conducting model and parameter recovery analyses on a held-out data set (described below). Prior to this step, we determined priors from independent (held-out) behavioural data. Importantly, determination of priors, parameter/model recovery analyses and final model specification were performed before seeing the data used in the main analysis.

In the HGF, belief updates at each hierarchical level *i* have the following general form:

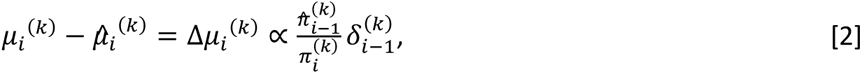

where the hat operator is used to indicate predictions. The term *δ_i_*_-1_ represents the PE about the hidden state one level below and is weighted by a ratio of the precisions (i.e., inverse variances) of the prediction about the level below and the belief at the current level. Thus, these belief updates are pwPEs. In this paper and our analysis, we denote the pwPE about skin temperature outcome by *ɛ*_2_. For further details, please refer to the supporting information.

### Perceptual model 2: Rescorla-Wagner model

The Rescorla-Wagner (RW) model (76) is a standard associative learning model with a fixed learning rate, *α*. Using the same notation as above, updates in this model take the following form:

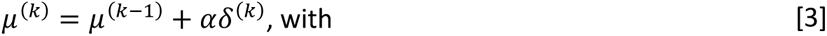

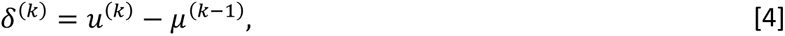

where *u*^(*k*)^ is the input at time point *k*, and the PE *δ* represents the difference between the input and the prediction from the previous time point.

### Estimation of priors from independent data

In order to define prior means and variances of free model parameters in an independent and empirical manner, we used our independent behavioural data set. For the 20 participants in this held-out data set, no MRI data were collected, but the same experimental procedure was followed in our behavioural laboratory. We fit each of the three models in our model space – 2-level HGF, 3-level HGF, and RW model – to participants’ behavioural responses, setting the initial prior means and variances to their default values in the HGF toolbox.

To ensure that the default prior parameter values represented reasonable *a priori* estimates for the STL task, we performed a qualitative analysis (described in the supporting information) and concluded that the default prior parameter settings in the HGF toolbox represented a reasonable choice for our independent data set. Given this result, we used the distribution of maximum a posteriori (MAP) point estimates across subjects in the held-out behavioural data set to specify priors for the model-based analysis of behaviour during fMRI in the analysis of interest. Specifically, we used the distribution of MAP estimates obtained by inverting the models (under default HGF priors) using the held-out (behavioural) data set and determined their means and variances by means of a fast minimum covariance determinant algorithm (77) that is robust to outliers. These means and variances served to define new priors (referred to below as “empirical priors”) that were used for model-based analysis of behaviour during the fMRI paradigm. Parameters that were not estimated were fixed to their default settings in the HGF toolbox. The resulting parameter settings for all candidate models are summarised in Table S7.

### Selection of HGF variant

To determine which Bayesian model (i.e., the 2- or 3-level HGF) to include in the final model space for the main (fMRI) data set, we performed model and parameter recovery analyses on the held-out data set (see previous section). We simulated responses for 100 synthetic subjects with each of the three models in our initial model space (i.e., a total of 300 simulated response trajectories), with values of the free parameters sampled from the empirical priors (see previous section). Each model was fit to each simulated trajectory, allowing us to compare the log model evidences (LME) across the three models and to identify a winning model for each simulated trajectory. A confusion matrix summarising the number of correctly identified models showed that the 2-level HGF was more easily recoverable in the simulations (see Figure S10). To assess the ability of each model to recover the true (data-generating) parameters, we compared the parameter estimates obtained by model inversion to the parameter values used to generate the response trajectories (see Figure S10). Based on the model and parameter recovery results, when applied to data from the STL task, the 2-level HGF was more easily distinguishable from the RW model than the 3-level HGF and its parameters were better recoverable. Therefore, we included the 2-level HGF and the RW model in the final model space for our analysis.

### Model comparison

To identify which of the two learning models (i.e., the 2-level HGF or the RW model) provides the more plausible explanation of participants’ trial-wise responses (and thereby address Hypothesis 1), we used random-effects Bayesian model selection (35, 36) (BMS). BMS compares estimates of log model evidences (LME; for each subject and each model) and describes the relative plausibility of different models in terms of posterior model probabilities and (protected) exceedance probabilities. The winning model was selected as the model with the highest protected exceedance probability (PEP), a measure that quantifies the probability that a particular model is more likely than all other models considered, corrected for the possibility that differences in model evidence are due to chance (36).

Posterior estimates of the free model parameters, computed from the MAP estimates obtained by inverting the models on each participant’s data, are listed in Table S8.

### Model-based regressors

The prediction and PE trajectories estimated with the winning model were used to generate subject-specific regressors of interest for the MRI analyses. Since the chosen model was the 2-level HGF, we modelled trajectories of pwPEs. Empirical evidence indicates that warm and cool temperatures reach the brain via separate ‘labelled lines’, with crosstalk between these pathways occurring at various stages of processing (reviewed in (78)). Therefore, we represented the computational quantities of interest using separate regressors for warm and cool temperatures.

To obtain prediction trajectories concerning skin temperature (T), we transformed the model-derived trajectory for 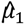 (estimated in contingency space) back into temperature space, according to the cue presented on each trial:

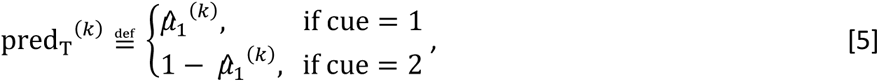

where pred_T_^(*k*)^ = 1 corresponds to a maximal prediction of skin warming and pred_T_^(*k*)^ = 0 to a maximal prediction of skin cooling.

To convert the model-derived trajectory for ε_2_ into temperature space, we took its absolute value:

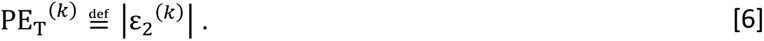

In the setting of our task, which entailed (i) categorical predictions of outcomes and (ii) coupled probabilities in the two learning trajectories, this gives equivalent results to modelling two separate learning processes for warm and cool temperature outcomes and computing the respective pwPE trajectories in temperature space. (For a detailed explanation, see the supplementary material of (12)).

### MRI analysis

For each participant, we specified a voxel-wise general linear model (GLM) including the following regressors: cue presentation on trials where the participant predicted skin warming and (as a separate regressor) parametric modulation by trial-wise prediction values; cue presentation on trials where the participant predicted skin cooling and parametric modulation by trial-wise prediction values; warm stimulus presentation and parametric modulation by trial-wise pwPE values; cool stimulus presentation and parametric modulation by trial-wise pwPE values; missed/late responses; and regressors representing head motion and physiological noise (see supporting information). The four parametric modulators were orthogonalised with respect to their corresponding main regressors. All regressors were convolved with the canonical haemodynamic response function (HRF).

We specified the following contrasts of interest: pwPE trajectories for warm and cool temperatures (as separate conditions), prediction trajectories for warm and cool temperatures (as separate conditions), average of warm and cool pwPEs, average of warm and cool predictions, the difference between warm and cool pwPEs, and the difference between warm and cool predictions. At the group level, we performed one-sample t-tests on the first-level contrast images in a random-effects GLM.

To test the hypothesis that activity in the insula is correlated with interoceptive predictions and (pw)PEs (Hypothesis 2), respectively, we used a mask of the bilateral insula (based on the Brainnetome atlas (79), as pre-specified in our analysis plan). To test the hypothesis that dopaminergic midbrain activity is correlated with interoceptive predictions and (pw)PEs (Hypothesis 3), we performed an analysis of pre-specified ROIs, i.e. the substantia nigra (SN) and ventral tegmental area (VTA), using the same anatomical mask as Iglesias et al. (12), which was derived from magnetisation transfer-weighted structural MRI images (80). We applied peak-level family-wise error (FWE) correction for multiple comparisons (*p* < 0.05) using Gaussian random field theory (81). We further adjusted significance thresholds because we employed two one-tailed t-tests in SPM (to test for both directions of effects), and for the number of ROIs (2). Therefore, significance corresponded to *p* < 0.01 (rounded to 2 decimal places) at the peak level (FWE-corrected).

The ROI analyses were followed by a whole-brain analysis to test the hypothesis that interoceptive predictions and (pw)PEs are reflected in the activity of large-scale networks distributed across the brain (Hypothesis 4). Adjusting for the use of two one-tailed t-tests, we defined significance as *p* < 0.025 at the cluster level (FWE-corrected) with a cluster-defining threshold of *p* < 0.001. To provide an indication of the locations of significant activation clusters, we identified the locations of cluster peaks (with the exception of the insula; see next section) using the Brainnetome atlas (79).

A summary of all statistical tests is presented in Table S1.

### Deviations from the preregistered analysis plan

We had specified that we would report effect sizes (Cohen’s *d* with 95 % confidence intervals) for activation peaks in all our analyses. However, since estimating effect sizes in neuroimaging is problematic (82), we decided to abandon this step.

Further, we did not identify activation peaks in the insula using the Brainnetome atlas (79), as indicated in our analysis plan. The reason for this deviation is that insular labels in the Brainnetome atlas refer to cytoarchitectonic subregions, but the underlying parcellation scheme is based on connectivity information from neuroimaging studies. Moreover, recent parcellations of the insula (e.g. (79, 83–85)) differ from each other. Since theories of interoceptive Bayesian inference make specific proposals related to the cytoarchitecture of insular subregions (e.g. (20, 22)), we chose to refrain from using labels that could lead to misinterpretation or confusion (e.g., the region referred to as the dorsal agranular insula in the Brainnetome atlas is labelled as dysgranular in other atlases (83, 85)). Instead, we decided to use broad topographic labels only (e.g. dorsal anterior insula).

### Code review

An independent code review was performed by one of the coauthors (SI) on the full analysis pipeline. The analysis code is available on the ETH GitLab repository (https://gitlab.ethz.ch/tnu/code/toussaintetal_compi_stl).

