## Supporting information for "Thermoceptive predictions and prediction errors in the anterior insula"

### Results

*Whole-brain analysis: post-hoc findings*

In a post-hoc analysis (which was not included in our pre-specified analysis plan), we investigated in which brain regions activity was correlated with temperature changes. As our experiment design did not include an explicit thermal baseline condition, we tested for the difference between warm and cool temperatures. Significant activations with the contrast warm < cool were observed in five clusters containing (inter alia) the bilateral (mainly right mid-posterior) insula, primary somatosensory and motor cortices, the left thalamus, and the right orbitofrontal cortex (see Figure S6 and Table S6). No significant clusters were observed with the contrast warm > cool.

### Methods

In the following sections, we expand on the data collection and analysis methods described in the main text with additional details.

*Pre-screening*

All participants were pre-screened online to ensure they did not display clinically significant levels of depression or central sensitization syndrome, a group of medically indistinct disorders such as fibromyalgia, chronic fatigue, and irritable bowel syndrome (see main text). Further exclusion criteria were defined as past or present mental or somatic health conditions, medication intake within 7 days prior to the study, past or present drug abuse (except nicotine), dermatological conditions involving the forearm, and pregnancy. Participation in a similar learning study or a neural stimulation study resulted in a temporary exclusion of 7 days. Volunteers for the fMRI study were pre-screened for MRI-compatibility.

*Sample size*

Since we tested a novel experimental paradigm, we did not have reliable effect size estimates. In our analysis plan, we had specified a Sequential Bayes Factor (1) (SBF) design with regard to the behavioural data: a first evaluation of the group Bayes Factor (2) (GBF) for the alternative models specified by Hypothesis 1 (see Table S1) was planned after N = 50 participants, followed by subsequent evaluations of the GBF with each additional data set, up to a resources-driven maximum sample size of 100 participants. As we used random-effects Bayesian model selection, we additionally required a protected exceedance probability > 0.9 (see *Model comparison*). However, due to technical issues with the thermostimulation system, data collection was repeatedly interrupted. Since data acquisition for our study was limited by time constraints (end of grant funding), we were forced to abandon our predefined SBF design and instead collected as many data sets as possible within the available time, resulting in a final sample of 44 data sets. At this point, the GBF in favour of the winning model (see below) exceeded our pre-defined stopping threshold of 20, whereas the protected exceedance probability of 0.77 fell short of our second stopping criterion. For the estimation of priors in the computational models, we collected an independent behavioural data set, for which we had pre-specified a fixed sample size of N = 20 in our analysis plan.

*Experimental protocol*

As described in the main text, the experiment began with a 5-minute adaptation period in which a continuous stimulation at the baseline temperature (32 °C) was applied to ensure that the skin in contact with the thermode adapted completely to this temperature. Next, we familiarised the participant with the experimental stimuli and performed several control checks. These steps are described in detail below:

We presented a series of 5 consecutive cool (27 °C) and 5 consecutive warm (39 °C) stimuli, along with on-screen instructions, in which the cool and warm stimulus sequences were explicitly described as cooling and warming the skin. We then presented the same series of 5 cool and 5 warm stimuli again. The participant was asked to (verbally) describe their perception of temperature and any other sensations. The participant's descriptions were documented in an electronic database (REDCap electronic data capture tools hosted at ETH Zurich (3, 4)). Following this introduction, we presented a sequence of 5 warm, 5 cool and 5 thermoneutral (i.e., baseline temperature = 32 °C) stimuli in pseudo-random order and asked the participant to identify each stimulation as warming, cooling, or maintaining neutral skin temperature using a response box. If the participant misclassified more than 4 stimulations (corresponding to less than 73.3 % correct), this step was repeated. If more than 4 stimulations were misclassified a second time, the participant was excluded from the experiment.

The participant then performed the skin temperature learning (STL) task, as described in the main text. After completion of the STL task, the participant was asked to identify 15 stimulations (5 warm, 5 cool and 5 thermoneutral) in a different pseudo-random sequence. This step provided an indication whether the participant could still distinguish warm and cool temperatures after the 30-minute task, allowing us to rule out complete habituation. Finally, we performed a 5-minute structural scan. The participant completed a debriefing questionnaire outside the scanner.

*Criteria for exclusion and replacement of participants*
As described in the main text, prior to data collection, we had defined criteria in our analysis plan for excluding and replacing participants due to data issues. These criteria in the categories of thermosensation, behavioural responses, and MRI data quality are further described below.

(i) Thermosensation
Thermal stimuli delivered at a fixed location can give rise to adaptation effects in sensory afferents, or habituation effects in the central nervous system (5). To mitigate the possibility of sensory adaptation, we selected stimulus temperatures well above and below human hot and cold detection thresholds (as verified in a thermostimulation pilot study involving only behavioural measurements; N = 20). The warm and cool temperatures in our experiment lie well outside the zone of physiological zero; therefore, complete adaptation to these temperatures was not expected to take place (6). To mitigate the possibility of central habituation, each temperature pulse was immediately followed by a much longer stimulation at the thermoneutral baseline temperature. As a control check for habituation, we presented short sequences of warm, cool and thermoneutral stimuli (in pseudo-random order) before and after the task. These sequences allowed us to verify whether participants could still distinguish warming and cooling of their skin after repeated stimulation, and thereby to infer if complete habituation had occurred. Participants who misidentified 1) more than 4 stimulations, or 2) more than 3 stimulations from a single stimulus category (i.e., warm, cool or thermoneutral) in the control sequence following the skin temperature learning (STL) task were excluded and replaced. This resulted in the replacement of 3 participants in the MRI data set and 2 participants in the behavioural data set.

We further implemented control checks to ensure that our results reflect the neural processing of innocuous thermal stimuli and are largely free of confounding perceptual effects. Some healthy individuals experience non-painful burning or stinging during skin cooling to mild temperatures, a phenomenon termed paradoxical heat sensation (7–9), which appears to activate a region of the insula that is suppressed during normal cool perception (10). Therefore, we probed participants for reports of paradoxical heat in the debriefing questionnaire, an approach we had tested in the pilot study mentioned above. Participants who reported that the cool stimulation did not feel cool, but elicited a burning or stinging sensation, were excluded and replaced. We replaced 3 participants in the MRI data set based on this criterion. No participants in the behavioural sample reported paradoxical heat sensations.

Finally, we probed participants for reports of pain during the presentation of the thermal stimulations at the beginning of the experiment, and in the debriefing questionnaire at the end of the experiment. No participants reported pain.

(ii) Behavioural responses
We computed the percentage of each participant's correct predictions and missed responses. Participants who performed below chance level (i.e., less than 50 % correct predictions) or failed to respond on 15 % of trials or more (> 22 missed responses) were excluded and replaced. Based on these criteria, 2 and 1 participant(s) were replaced in the MRI and behavioural data sets, respectively.

(iii) MRI data quality
All MRI data sets were inspected (visually, using the TSDiffAna toolbox; https://www.fil.ion.ucl.ac.uk/spm/ext/#TSDiffAna and SPM12; http://www.fil.ion.ucl.ac.uk/spm) to ensure that preprocessing was successful and that there was no signal dropout in the insula or the dopaminergic midbrain. In one data set, we detected large distortions in several volumes. Prior to preprocessing, we replaced 172 affected volumes in this data set by linearly interpolating between the nearest confound-free neighbouring volumes. In the fMRI analyses, we included a stick regressor for each of the replaced volumes in the general linear model (GLM) for this participant. The total number of stick regressors included for this participant was below the pre-specified threshold (based on subject-level degrees of freedom) described in the next paragraph.

We controlled for correlations of task and motion in the STL task design by counterbalancing the order of responses (left warm – right cool/ left cool – right warm) across participants, and by jittering SOA lengths. Further, we pre-specified the exclusion of participants whose data were excessively affected by abrupt movements. To this end, our analysis plan defined a threshold for the maximum tolerable number of motion confound regressors (see **Error! Reference source not found.**) such that their inclusion in the GLM would reduce the subject-level degrees of freedom by no more than 20 % of the total number of volumes. No data sets were excluded due to excessive motion confounds.

*Preprocessing*
In our statistical analyses, we accounted for head motion and physiological confounds by including two types of nuisance regressors. Head motion was accounted for by the six translation and rotation parameters resulting from image realignment, and their temporal derivatives. Further, we computed framewise displacement (11) (FD) using the implementation in the PhysIO toolbox (12) in the TAPAS software (13) (https://www.tnu.ethz.ch/en/software/tapas), and included a stick regressor for each volume with an FD greater than 0.9 mm (i.e., motion censoring). We corrected for physiological BOLD signal confounds by generating physiological noise regressors using measured respiratory and pulse oximetry data in the PhysIO toolbox. Specifically, we created RETROICOR regressors (14) with Fourier expansions of physiological phases based on Harvey et al.'s recommendations (15) (i.e., $3^{rd}$-order, $4^{th}$-order, and $1^{st}$-order expansions for cardiac, respiratory and interaction models, respectively). We also created regressors to account for slow respiratory and heart rate fluctuations. The respiratory and cardiac responses were modelled by convolving these physiological signals with response functions to respiratory volume per unit time (16) (RVT) and heart rate variability (17) (HRV).

*Computational modelling of behaviour: the Hierarchical Gaussian Filter*
Below, we provide a brief outline of the Hierarchical Gaussian Filter (HGF) implementation in our study. A detailed description of the inversion scheme and the specific update equations can be found in the papers that introduced the HGF (9, 36).

In our three-level HGF implementation, the first level represents beliefs about environmental states $x_1$ (corresponding to the binary cue-outcome association, for example "1" represents a warm stimulus following cue 1 and "0" represents a cool stimulus following cue 1), the second level represents beliefs about the tendency $x_2$ of cue 1 to predict a warm stimulus, and the third level represents beliefs about the environmental volatility $x_3$ (i.e. the degree to which the tendency that cue 1 predicts a warm stimulus changes over trials). At levels 2 and 3, the hidden states $x_i$

evolve as a Gaussian random walk with mean $\mu_i$ and variance $\pi_i^{-1}$, where the latter depends on the state at level $i + 1$. At the first level, the states in $x_2$ are mapped onto cue-outcome associations via a sigmoid transformation. A graphical representation of this model is shown in Figure S8.

As new sensory information arrives, the agent updates his/her beliefs about these hidden states. These belief updates have the following general form:

$$\mu_i^{(k)} - \hat{\mu}_i^{(k)} = \Delta\mu_i^{(k)} \propto \frac{\hat{\pi}_{i-1}^{(k)}}{\pi_i^{(k)}} \delta_{i-1}^{(k)}, \quad [1]$$

where the hat operator is used to indicate predictions. The term $\delta_{i-1}$ represents the PE about the hidden state one level below and is weighted by a ratio of the precisions (i.e., inverse variances) of the prediction about the level below and the belief at the current level. Thus, these belief updates are pwPEs. In our three-level implementation, we denote the pwPE about skin temperature outcome (that is used to update the estimate of $x_2$) by $\varepsilon_2$, and the pwPE about cue-outcome contingency (that is used to update the estimate of $x_3$) by $\varepsilon_3$.

As depicted in Figure S8, the model contains parameters $\kappa$ (the coupling strength between the second and third levels), $\omega_2$ (the constant component of volatility on the second level), and $\omega_3$ (how fast learning about environmental volatility on the third level occurs). These parameters can be estimated for each participant to account for individual differences in learning. We modelled $\omega_2$ and $\omega_3$ as free parameters, while holding $\kappa$ fixed. In our two-level HGF implementation, the third level was removed from the computational hierarchy described above. The two-level HGF is thus a simpler Bayesian model with only one free parameter, $\omega_2$ (the tonic volatility on the second level), that still allows for a dynamic learning rate and takes the precision of beliefs into account. It follows that in this model variant pwPEs only occur at the second level (denoted by $\varepsilon_2$).

*Estimation of priors from independent data*

In order to define prior means and variances of free model parameters in an independent and empirical manner, we used our independent behavioural data set. For the 20 participants in this held-out data set, no MRI data were collected, but the same experimental procedure was followed in our behavioural laboratory. We fit each of the three models in our model space – 2-level HGF, 3-level HGF, and RW model – to participants' behavioural responses, setting the initial prior means and variances to their default values in the HGF toolbox.

To ensure that the default prior parameter values represented reasonable *a priori* estimates for the STL task, we performed a qualitative analysis, in which we examined the distributions of perceptual model belief trajectories at the outcome level of 100 synthetic subjects (see Figure S9). These prior densities were created by randomly sampling values of the free parameter(s) in each perceptual model from their respective prior distributions. (Note that this is analogous to an analysis of the prior predictive density of each model. However, strictly speaking, the plots in Figure S9 do not display the typical prior predictive densities of generative models of behaviour, as we analysed the outcome level of the perceptual models, rather than synthetic behavioural responses.) As seen in Figure S9, the initial prior settings allowed each model to explain a wide range of plausible trajectories and did not produce many implausible trajectories. This last point was desirable given our pre-specified outcome-neutral criteria, i.e., the exclusion of participants who failed to perform the task correctly or did not learn anything about the underlying probabilistic structure. Based on this qualitative analysis, we concluded that the default prior parameter settings in the HGF toolbox represented a reasonable choice for our independent data set.

Given this result, we used the distribution of maximum a posteriori (MAP) point estimates across subjects in the held-out behavioural data set to specify priors for the model-based analysis of behaviour during fMRI in the analysis of interest. Specifically, we used the distribution of MAP estimates obtained by inverting the models (under default HGF priors) using the held-out (behavioural) data set and determined their means and variances by means of a fast minimum covariance determinant algorithm (76) that is robust to outliers. These means and variances served to define new priors (referred to below as “empirical priors”) that were used for model-based analysis of behaviour during the fMRI paradigm. Parameters that were not estimated were fixed to their default settings in the HGF toolbox. The resulting parameter settings for all candidate models are summarised in Table S7.

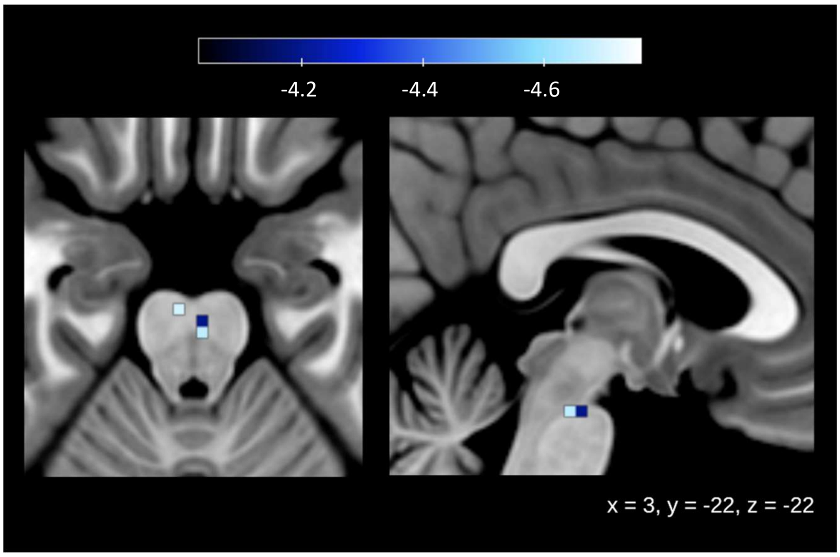


**Fig. S1.** Midbrain responses to predictions. Deactivations associated with predictions about skin temperature outcome, $\hat{\boldsymbol{\mu}}_{\mathbf{1}}$, shown at a threshold of p < 0.01, peak-level FWE-corrected for multiple comparisons in the SN/VTA.

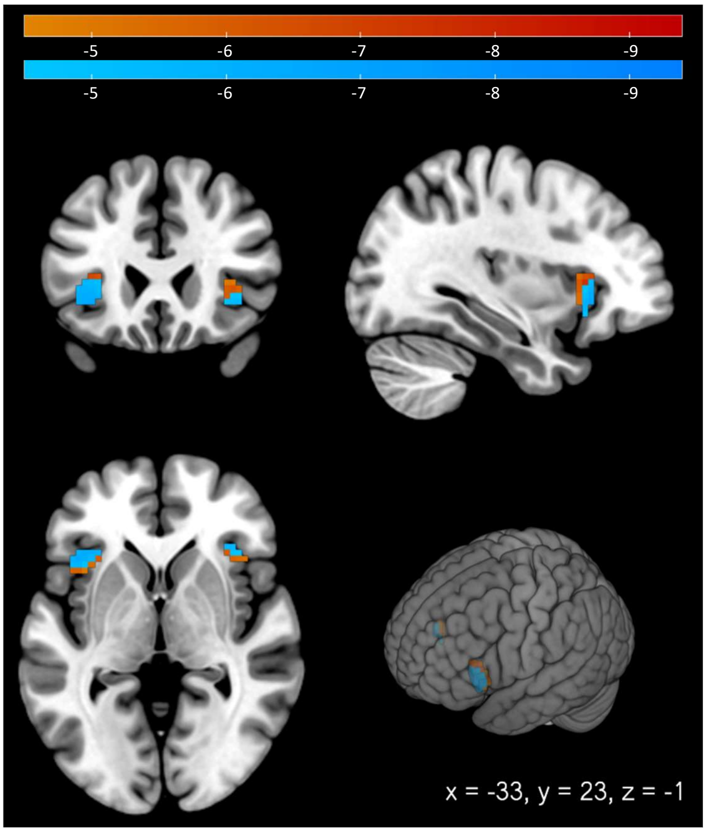


**Fig. S2.** Insula analysis: findings of predictions about skin warming and cooling. Deactivations associated with predictions ($\hat{\mu}_1$) about skin warming (in orange/red) and cooling (in blue) shown at a threshold of $p$ < 0.01, peak-level FWE-corrected for multiple comparisons in the bilateral insula.

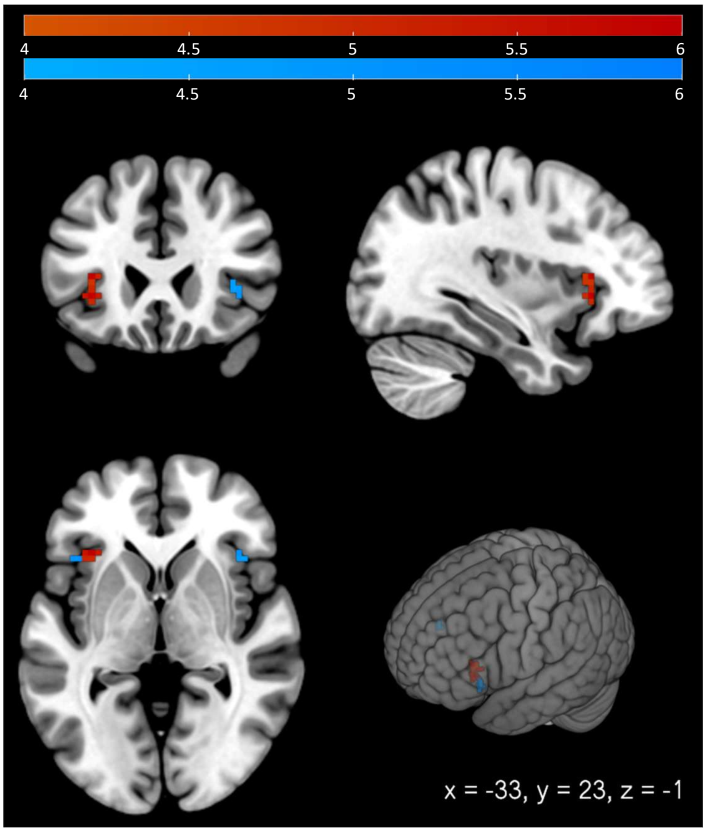


**Fig. S3.** Insula analysis: findings of precision-weighted prediction errors about skin warming and cooling. Activations associated with precision-weighted prediction errors ($\varepsilon_2$) about skin warming (in orange/red) and cooling (in blue) shown at a threshold of $p < 0.01$, peak-level FWE-corrected for multiple comparisons in the bilateral insula.

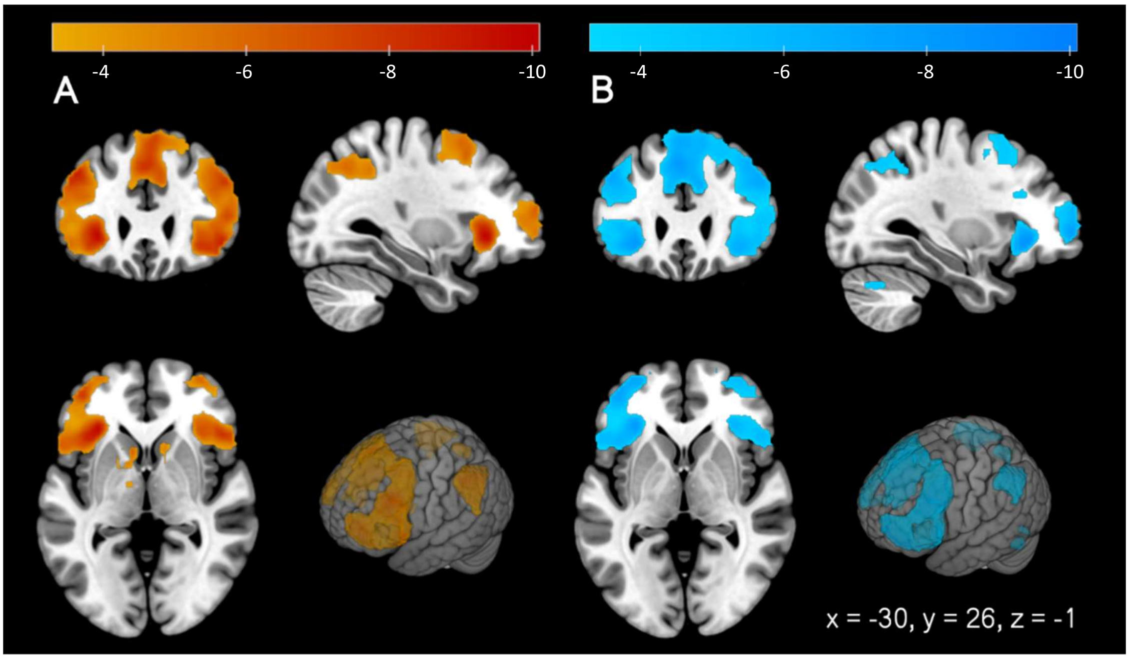


**Fig. S4.** Whole-brain findings of predictions about skin warming and cooling. Deactivations associated with predictions ($\hat{\mu}_1$) about skin warming (in orange/red) and cooling (in blue) shown at a cluster-level threshold of $p < 0.025$, FWE-corrected for multiple comparisons across the whole brain (cluster-defining threshold of $p < 0.001$).

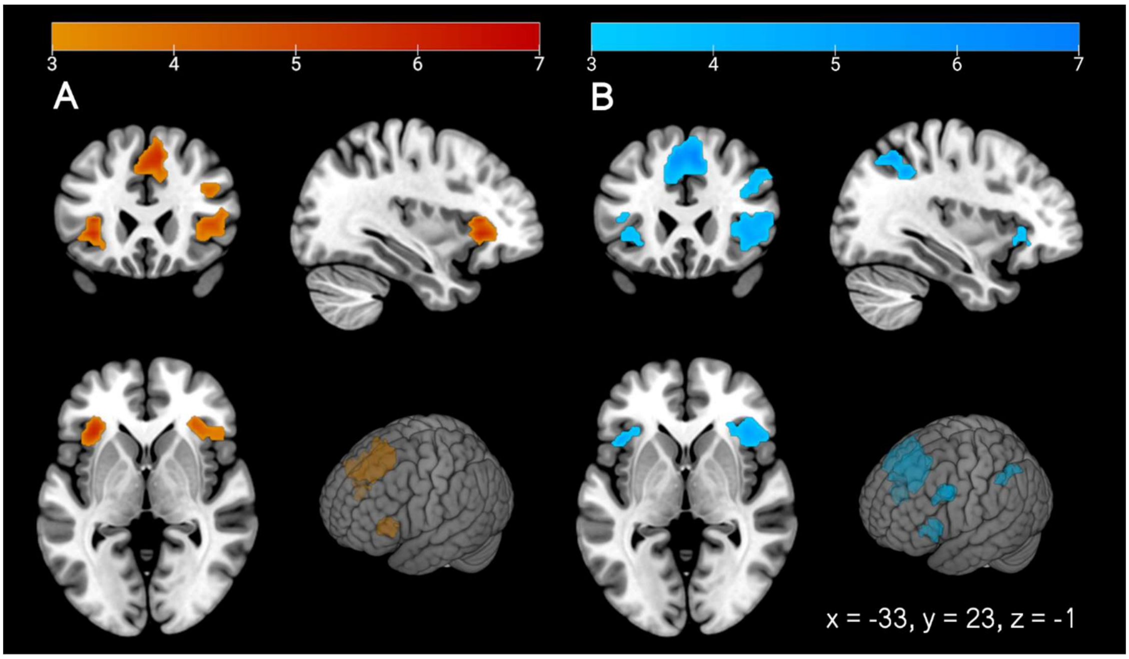


**Fig. S5.** Whole-brain findings of precision-weighted prediction errors about skin warming and cooling. Activations associated with precision-weighted prediction errors ($\varepsilon_2$) about skin warming (in orange/red) and cooling (in blue) shown at a cluster-level threshold of $p < 0.025$, FWE-corrected for multiple comparisons across the whole brain (cluster-defining threshold: $p < 0.001$).

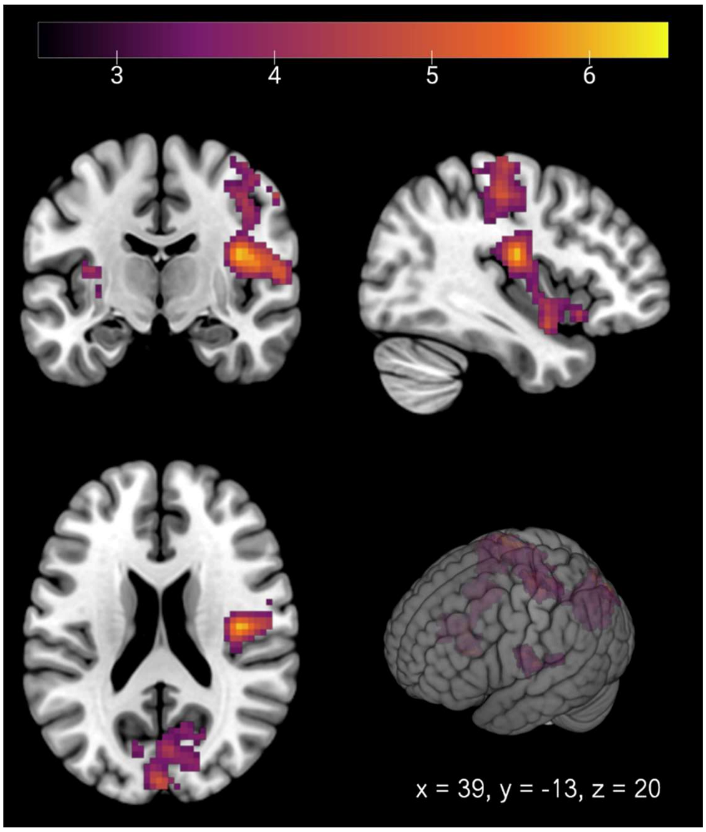


**Fig. S6.** Whole-brain analysis of the difference between warm and cool temperatures. Activations associated with the contrast warm < cool shown at a cluster-level threshold of $p < 0.025$, FWE-corrected for multiple comparisons across the whole brain (cluster-defining threshold: $p < 0.001$).

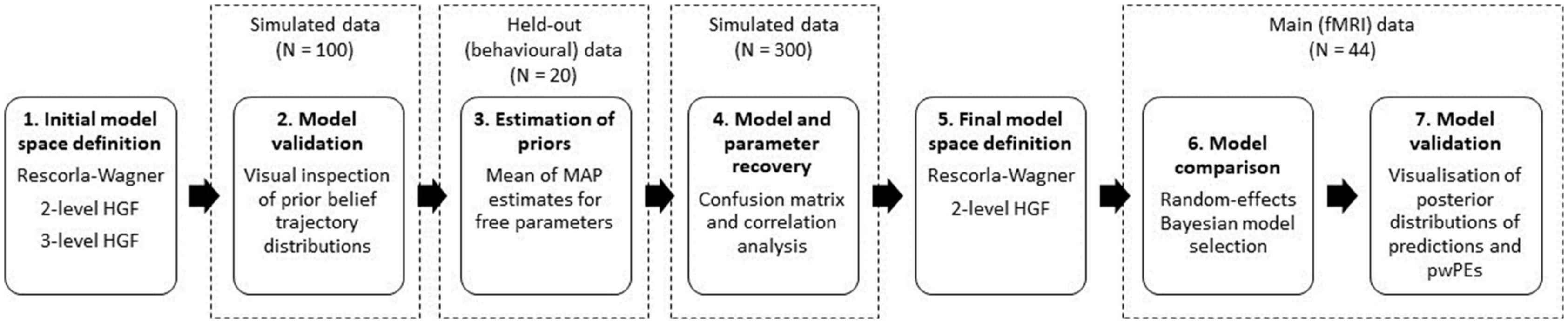


**Fig. S7.** Pipeline for computational modelling of behaviour in the skin temperature learning (STL) task.

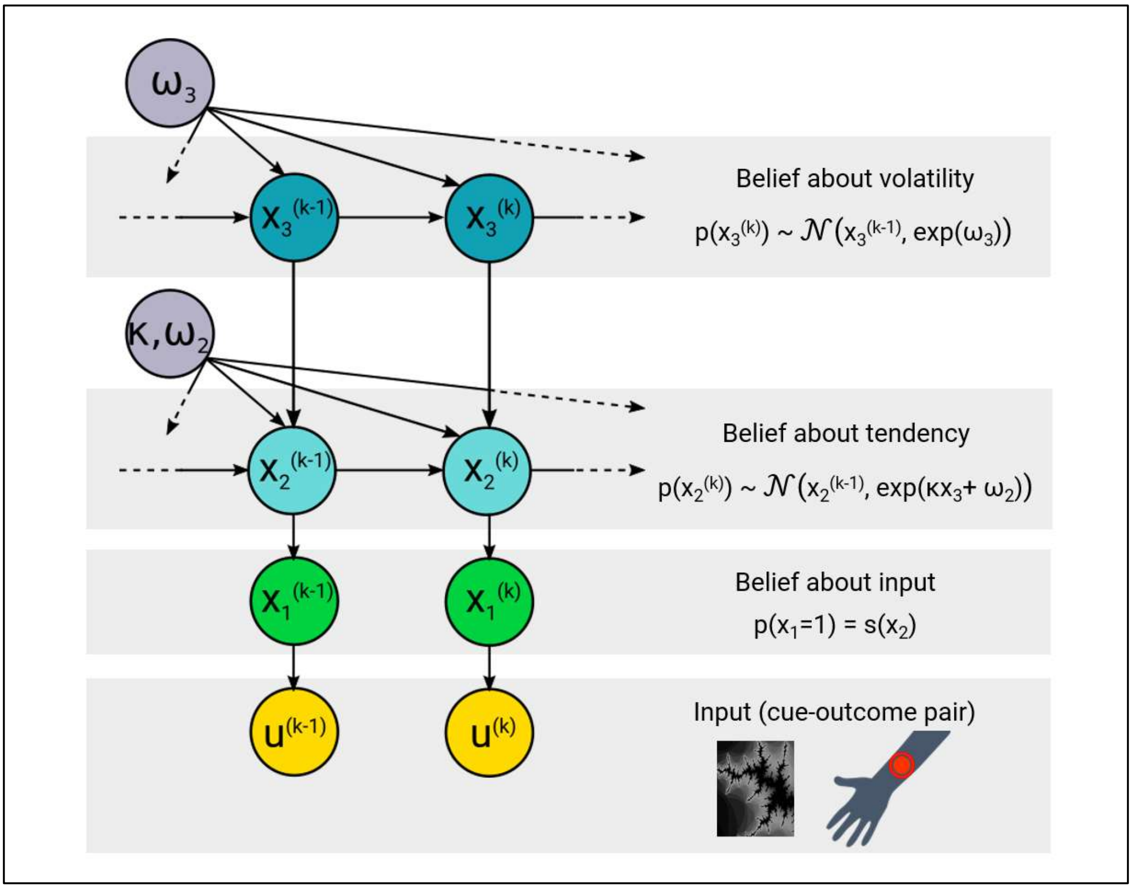


**Fig. S8.** Hierarchical Gaussian Filter for modelling the skin temperature learning (STL) task – the three-level Hierarchical Gaussian Filter (HGF3). The belief $x_1$ represents the cue outcome-association, $x_2$ the tendency of the cue-outcome association, and $x_3$ the environmental volatility (i.e., the degree to which $x_2$ changes over time). Each of these beliefs is parameterised by a mean $\mu$ and variance $\sigma$.

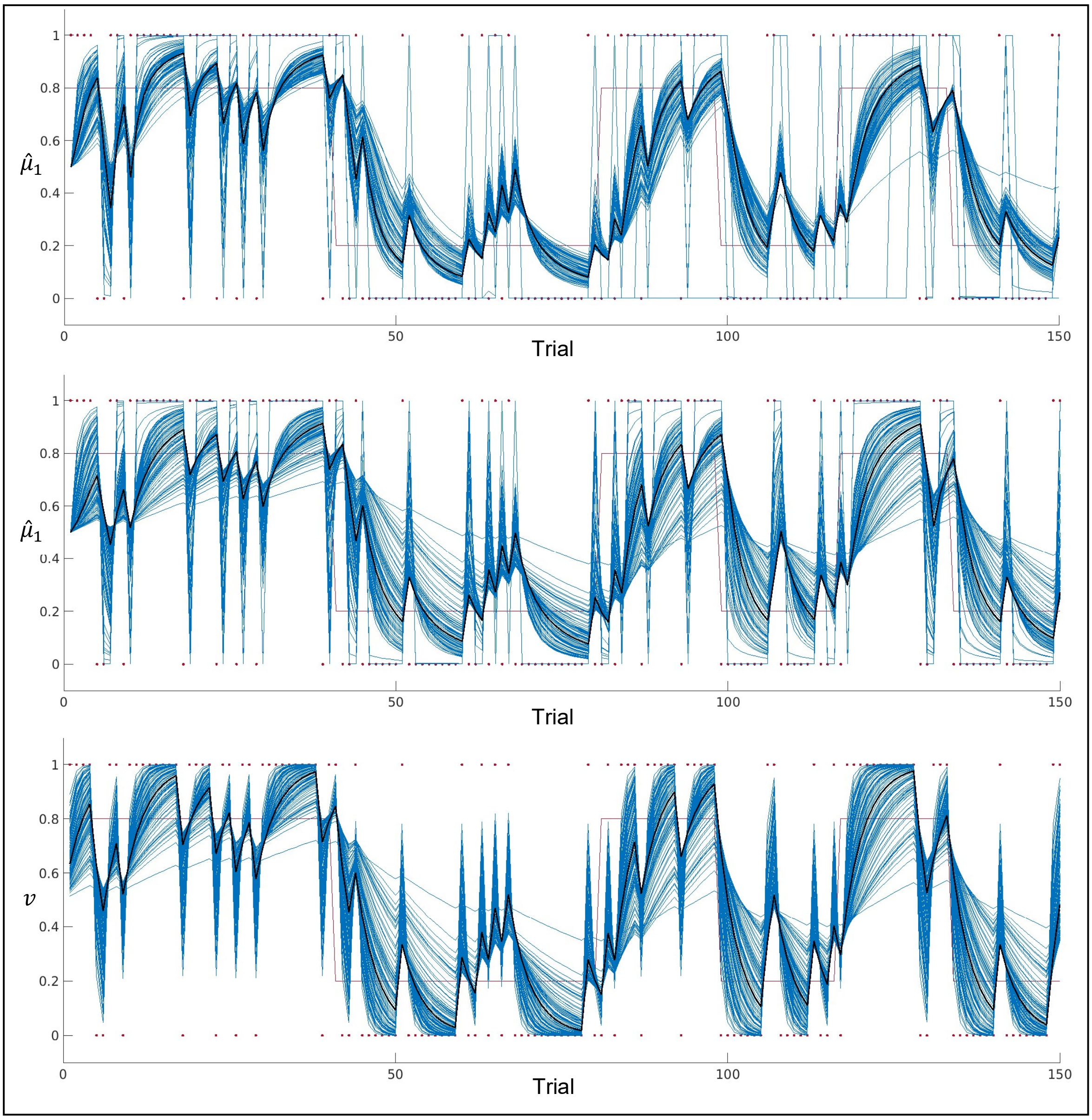


**Fig. S9.** Prior densities of the perceptual models. *Top:* Belief trajectories at the outcome level of the three-level HGF ($\hat{\mu}_1$), generated using 100 randomly sampled values of the free perceptual model parameters ($\omega_2$ and $\omega_3$) from their respective prior densities. The thick black trajectory was generated by setting the free parameters to their prior means. Red dots represent the sequence of binary inputs and the red line represents the underlying probabilistic structure of the task. *Middle:* Belief trajectories at the outcome level of the two-level HGF ($\hat{\mu}_1$), generated using 100 randomly sampled values of the free perceptual model parameter ($\omega_2$) from its prior density. The thick black trajectory was generated by setting the free parameter to its prior mean. Red dots represent the sequence of binary inputs and the red line represents the underlying probabilistic structure of the task. *Bottom:* Belief trajectories at the outcome level of the Rescorla-Wagner model ($v$), generated using 100 randomly sampled values of the free perceptual model parameter ($\alpha$) from its prior density. The thick black trajectory was generated by setting the free parameter to its prior mean. Red dots represent the sequence of binary inputs and the red line represents the underlying probabilistic structure of the task.

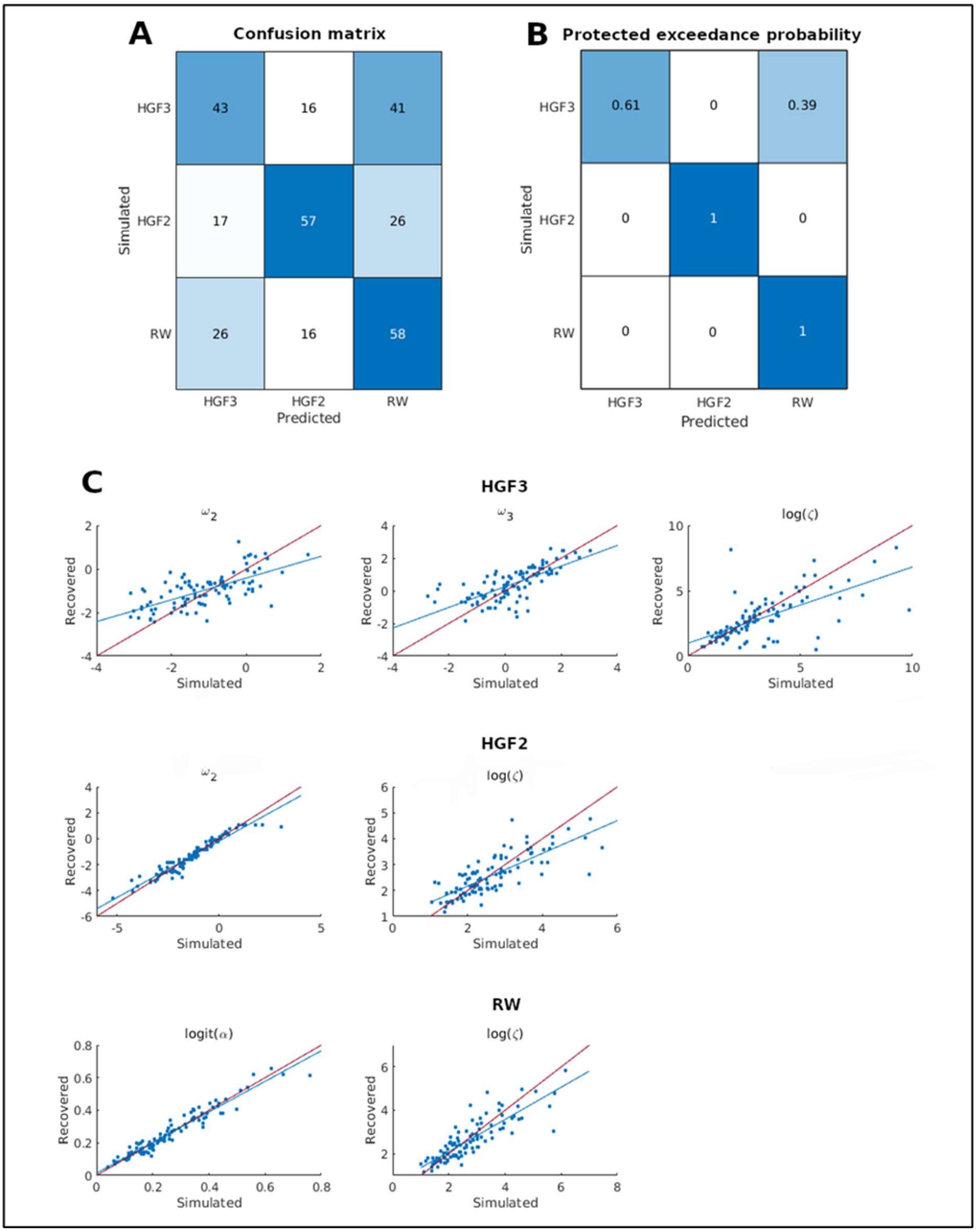


**Fig. S10.** Prior model and parameter recovery analyses. We considered 3 perceptual models in our prior model space: 3- and 2-level Hierarchical Gaussian Filters (HGF3 and HGF2, respectively), and a Rescorla-Wagner (RW) model. We fit each model to behavioural responses of participants in the held-out data set, and computed the resulting group means and variances for free parameters (see **Table S8**). We drew samples from these empirical parameter estimates to simulate responses for 100 synthetic subjects with each model. Each model was fit to each simulated trajectory, and the resulting log model evidences were compared across models based on a confusion matrix summarising the number of correctly identified models (A) and protected exceedance probabilities (B). The ability of each model to recover the data-generating parameters is shown in C. The line of best fit is indicated in blue, and the identity line in red.

**Table S1.** Design table.

| Question | Hypothesis | Analysis Plan | Interpretation given to different outcomes |
|---|---|---|---|
| 1. Which learning model best describes the learning of associations between predictive cues and physiological outcomes in a volatile environment? | A Bayesian learning model provides a better account of explicit predictions of skin temperature changes than a classical conditioning model. | Two computational models fit to individual behavioural data; random-effects Bayesian model selection at the group level. | PEP indicates the model with the highest explanatory power at the group level. |
| 2. Does neural activity in the insula reflect computational signatures of interoceptive predictive coding? | Interoceptive predictions and (precision-weighted) PEs are correlated with BOLD responses in the insula. | ROI fMRI analysis; group-level GLM: t-tests applied to contrast images ($PE_{warm} > 0$, $PE_{warm} < 0$, $PE_{cool} > 0$, $PE_{cool} < 0$, $predictions_{warm} > 0$, $predictions_{warm} < 0$, $predictions_{cool} > 0$, $predictions_{cool} < 0$) in the bilateral insula; FWE correction for multiple comparisons within the ROI. | Reject $H_0$ if $p < 0.01$* at the peak level (FWE-corrected); effect size measured using Cohen's *d* with 95 % CI |
| 3. Does neural activity in the dopaminergic midbrain reflect interoceptive prediction and PE signals? | Interoceptive predictions and (precision-weighted) PEs are reflected in BOLD responses in the dopaminergic midbrain. | ROI fMRI analysis; group-level GLM: t-tests applied to contrast images ($PE_{warm} > 0$, $PE_{warm} < 0$, $PE_{cool} > 0$, $PE_{cool} < 0$, $predictions_{warm} > 0$, $predictions_{warm} < 0$, $predictions_{cool} > 0$, $predictions_{cool} < 0$) in the dopaminergic midbrain; FWE correction for multiple comparisons within the ROI. | Reject $H_0$ if $p < 0.01$* at the peak level (FWE-corrected); effect size measured using Cohen's *d* with 95 % CI |
| 4. What is the extent of the brain network involved in processing predictions and PEs about skin temperature changes? | Interoceptive predictions and (precision-weighted) PEs are correlated with BOLD responses in a large-scale brain network including anterior and mid-cingulate cortex; anterior, mid-, and posterior insular cortex; and the amygdala. | Whole-brain fMRI analysis; group-level GLM: t-tests applied to contrast images ($PE_{warm} > 0$, $PE_{warm} < 0$, $PE_{cool} > 0$, $PE_{cool} < 0$, $predictions_{warm} > 0$, $predictions_{warm} < 0$, $predictions_{cool} > 0$, $predictions_{cool} < 0$) in the whole brain; cluster-level FWE correction for multiple comparisons. | Reject $H_0$ if $p < 0.025$* at the cluster level (FWE-corrected) with a cluster-defining threshold of $p < 0.001$; effect sizes measured at the peak activation within the cluster using Cohen's *d* with 95 % CI |

*PE = prediction error; FWE = family-wise error; CI = confidence interval; ROI = region of interest; PEP = protected exceedance probability.*

** For questions 2-4 we specified α = 0.05. p-Values were (i) adjusted for testing nondirectional hypotheses using two one-tailed t-tests, and (ii) in questions 2-3, corrected for the number of ROIs.*

**Table S2.** Insula analysis: findings of predictions about skin warming and cooling. MNI coordinates and t-values for deactivations associated with predictions, $\hat{\mu}_1$.

| Predictions about warming | | | | | |
|---|---|---|---|---|---|
| | Hemisphere | x | y | z | t-score |
| Dorsal anterior insula | L | -33 | 23 | -1 | -9.39 |
| Dorsal anterior insula | R | 36 | 26 | -4 | -7.30 |
| Dorsal anterior insula | R | 45 | 20 | -7 | -5.37 |
| **Predictions about cooling** | | | | | |
| Dorsal anterior insula | L | -33 | 23 | -4 | -6.66 |
| Dorsal anterior insula | R | 36 | 26 | -4 | -6.04 |
| Ventral anterior insula | R | 39 | 20 | -13 | -4.59 |

All results: $p < 0.01$ peak-level FWE-corrected for multiple comparisons in the bilateral insula.

**Table S3.** Insula analysis: findings of precision-weighted prediction errors about skin warming and cooling. MNI coordinates and t-values for activations associated with precision-weighted prediction errors, $\varepsilon_2$.

| **Precision-weighted prediction errors about warming** | | | | | |
|---|---|---|---|---|---|
| | Hemisphere | x | y | z | t-score |
| Dorsal anterior insula | L | -33 | 23 | -1 | 6.08 |
| Dorsal anterior insula | L | -33 | 23 | 8 | 5.88 |
| **Precision-weighted prediction errors about cooling** | | | | | |
| Dorsal anterior insula | L | -39 | 20 | -7 | 5.93 |
| Dorsal anterior insula | R | 39 | 23 | -1 | 5.27 |

All results: $p < 0.01$ peak-level FWE-corrected for multiple comparisons in the bilateral insula.

**Table S4.** Whole-brain findings of predictions about skin warming and cooling. Extent of clusters (number of significant voxels) for deactivations associated with predictions ($\hat{\mu}_1$) about skin warming and cooling. MNI coordinates and t-values are listed for peaks within significant clusters. Deactivations for which $p < 0.025$ at the peak level are indicated with an asterisk.

| Extent | Anatomical label | Hemisphere | x | y | z | t-score |
|---|---|---|---|---|---|---|
| **Predictions about warming** | | | | | | |
| 4774 | Rostroventral area 8 | L | -42 | 23 | 38 | -10.10* |
| | Lateral area 12/47 | L | -30 | 26 | -1 | -9.97* |
| | Dorsomedial PFC | L | -3 | 20 | 44 | -9.82* |
| 692 | Caudal area 40 | L | -51 | -46 | 47 | -8.02* |
| | Rostrodorsal area 39 | L | -42 | -61 | 44 | -7.72* |
| | Rostrodorsal area 39 | L | -42 | -52 | 47 | -6.95* |
| 764 | Caudal area 40 | R | 51 | -46 | 41 | -7.24* |
| | Rostroventral area 39 | R | 57 | -52 | 38 | -7.03* |
| | Rostroventral area 39 | R | 48 | -55 | 41 | -6.72* |
| 206 | Ventral caudate | L | -12 | 14 | 2 | -7.19* |
| 112 | Precuneus | R | 9 | -61 | 47 | -5.41 |
| **Predictions about cooling** | | | | | | |
| 4942 | Lateral area 12/47 | L | -30 | 29 | -4 | -8.74* |
| | Dorsomedial PFC | L | -3 | 32 | 35 | -8.23* |
| | Middle frontal gyrus | L | -42 | 47 | -4 | -8.20* |
| 594 | Inferior parietal lobule/ Area 39 | R | 48 | -64 | 38 | -6.87* |
| | Rostrodorsal area 39 | R | 39 | -58 | 47 | -6.54* |
| 506 | Rostrodorsal area 39 | L | -42 | -52 | 47 | -6.54* |
| | Rostroventral area 39 | L | -45 | -67 | 44 | -6.48* |
| | Rostrodorsal area 39 | L | -36 | -58 | 44 | -6.14* |
| 85 | Cerebellum | L | -12 | -82 | -31 | -5.31 |
| 107 | Dorsomedial parietooccipital sulcus | R | 6 | -64 | 44 | -4.92 |

All clusters: $p < 0.025$, cluster-level FWE-corrected for multiple comparisons across the whole brain (cluster-defining threshold: $p < 0.001$). PFC: prefrontal cortex.

* $p < 0.025$, peak-level FWE-corrected for multiple comparisons across the whole brain.

**Table S5.** Whole-brain findings of precision-weighted prediction errors about skin warming and cooling. Extent of clusters (number of significant voxels) for activations associated with precision-weighted prediction errors ($\varepsilon_2$) about skin warming and cooling. MNI coordinates and t-values are listed for peaks within significant clusters. Activations for which $p < 0.025$ at the peak level are indicated with an asterisk.

| **Precision-weighted prediction errors about warming** | | | | | | |
|---|---|---|---|---|---|---|
| Extent | Anatomical label | Hemisphere | x | y | z | t-score |
| 340 | Dorsomedial PFC | L | -3 | 11 | 56 | 6.26* |
| | Dorsomedial PFC | R | 9 | 20 | 47 | 5.89* |
| | Dorsomedial PFC | L/R | 0 | 20 | 44 | 5.78* |
| 90 | Dorsal anterior insula | L | -33 | 23 | -1 | 6.06* |
| | Pars opercularis | L | -33 | 23 | 8 | 5.88* |
| 115 | Pars opercularis | R | 42 | 23 | 5 | 5.08* |
| 165 | Dorsal area 44 | R | 39 | 23 | 26 | 4.58 |
| **Precision-weighted prediction errors about cooling** | | | | | | |
| 426 | Dorsomedial PFC | R | 9 | 32 | 38 | 8.05* |
| | Dorsomedial PFC | R | 9 | 26 | 44 | 7.12* |
| | Dorsomedial PFC | L | -6 | 17 | 53 | 6.52* |
| 122 | Inferior frontal junction | L | -42 | 5 | 41 | 6.03* |
| 96 | Lateral area 12/47 | L | -39 | 20 | -7 | 5.92* |
| 183 | Caudal area 45 | R | 54 | 20 | 11 | 5.82* |
| 79 | Rostrodorsal area 49 | L | -36 | -55 | 44 | 5.72* |
| | Rostrodorsal area 39 | L | -36 | -49 | 38 | 5.58* |
| 247 | Inferior frontal junction | R | 48 | 14 | 44 | 4.89 |

All clusters: $p < 0.025$, cluster-level FWE-corrected for multiple comparisons across the whole brain (cluster-defining threshold: $p < 0.001$). PFC: prefrontal cortex.

* $p < 0.025$, peak-level FWE-corrected for multiple comparisons across the whole brain.

**Table S6.** Whole-brain analysis of the difference between warm and cool temperatures. Extent of clusters (number of significant voxels) for activations associated with the contrast warm < cool. MNI coordinates and t-values are listed for peaks within significant clusters. Activations for which $p < 0.025$ at the peak level are indicated with an asterisk.

| Extent | Anatomical label | Hemisphere | x | y | z | t-score |
|---|---|---|---|---|---|---|
| 1088 | Dorsal posterior insula | R | 39 | -13 | 20 | 6.11* |
| | Posterior orbitofrontal cortex | R | 24 | 5 | -16 | 5.76* |
| 478 | Medial superior occipital gyrus | L | -12 | -85 | 38 | 5.47 |
| 89 | Primary motor cortex (upper limb region) | L | -24 | -25 | 62 | 4.72 |
| 129 | Posterior parietal thalamus | L | -27 | -22 | -1 | 4.44 |
| 146 | Primary somatosensory cortex (lower limb region) | L | -15 | -34 | 47 | 4.40 |

All clusters: $p < 0.025$, cluster-level FWE-corrected for multiple comparisons across the whole brain (cluster-defining threshold: $p < 0.001$).

* $p < 0.025$, peak-level FWE-corrected for multiple comparisons across the whole brain.

**Table S7.** Parameter configurations and priors. Free parameters for each candidate model are indicated in bold; their prior means and variances were computed from the maximum a posteriori (MAP) estimates obtained by inverting the models on the held-out data set. All parameters have Gaussian priors. Prior means are given in native space, and prior variances in estimation (transformed) space.

| *3-level Hierarchical Gaussian Filter* | | | |
|---|---|---|---|
| Parameter | Prior mean | Prior variance | Transformation |
| $\mu_1^{(0)}$ | NaN | NaN | none |
| $\mu_2^{(0)}$ | 0 | 0 | none |
| $\mu_3^{(0)}$ | 1 | 0 | none |
| $\sigma_1^{(0)}$ | NaN | NaN | log |
| $\sigma_2^{(0)}$ | 0.1 | 0 | log |
| $\sigma_3^{(0)}$ | 0 | 0 | log |
| $\rho_1$ | NaN | NaN | none |
| $\rho_2$ | 0 | 0 | none |
| $\rho_3$ | 0 | 0 | none |
| $\kappa_1$ | 1 | 0 | log |
| $\kappa_2$* | 1 | 0 | log |
| $\boldsymbol{\omega_2}$ | **-1.07** | **1.00** | **none** |
| $\boldsymbol{\omega_3}$ | **0.30** | **1.21** | **none** |
| **Observation model $\boldsymbol{\zeta}$** | **2.59** | **0.29** | **log** |
| *2-level Hierarchical Gaussian Filter* | | | |
| Parameter | Prior mean | Prior variance | Transformation |
| $\mu_1^{(0)}$ | NaN | NaN | none |
| $\mu_2^{(0)}$ | 0 | 0 | none |
| $\mu_3^{(0)}$ | 1 | 0 | none |
| $\sigma_1^{(0)}$ | NaN | NaN | log |
| $\sigma_2^{(0)}$ | 0.1 | 0 | log |
| $\sigma_3^{(0)}$ | 0 | 0 | log |
| $\rho_1$ | NaN | NaN | none |
| $\rho_2$ | 0 | 0 | none |
| $\rho_3$ | 0 | 0 | none |
| $\kappa_1$ | 1 | 0 | log |
| $\kappa_2$* | 1 | 0 | log |
| $\boldsymbol{\omega_2}$ | **-1.33** | **2.36** | **none** |
| $\omega_3$ | $-\infty$ | 0 | none |
| **Observation model $\boldsymbol{\zeta}$** | **2.35** | **0.14** | **log** |
| *Rescorla-Wagner* | | | |
| Parameter | Prior mean | Prior variance | Transformation |
| $v^{(0)}$ | 0.5 | 0 | logit |
| $\boldsymbol{\alpha}$ | **0.24** | **0.67** | **logit** |
| **Observation model $\boldsymbol{\zeta}$** | **2.42** | **0.16** | **log** |

** Here $\kappa_2$ corresponds to $\kappa$ in* **Figure S8** *and the main text. The variable names listed here correspond to those in the HGF toolbox.*

**Table S8.** Posterior parameter estimates for candidate models in the final model space. Posterior means and variances of free parameters were computed from the maximum a posteriori (MAP) estimates obtained by inverting the models on the main data set. Posterior means and variances are both given in estimation (transformed) space.

| | *Hierarchical Gaussian Filter (2-level)* | |
|---|---|---|
| | Posterior mean | Posterior variance |
| $\omega_2$ | -1.58 | 0.44 |
| $\log(\zeta)$ | 0.76 | 0.22 |
| | *Rescorla-Wagner* | |
| | Posterior mean | Posterior variance |
| $\mathrm{logit}(\alpha)$ | -1.36 | 0.14 |
| $\log(\zeta)$ | 0.60 | 0.26 |
